# Landscape heterogeneity and soil biota are central to multi-taxa diversity for landscape restoration

**DOI:** 10.1101/2022.10.31.514517

**Authors:** Vannesa Montoya-Sánchez, Holger Kreft, Isabelle Arimond, Johannes Ballauff, Dirk Berkelmann, Fabian Brambach, Rolf Daniel, Ingo Grass, Jes Hines, Dirk Hölscher, Bambang Irawan, Alena Krause, Andrea Polle, Anton Potapov, Lena Sachsenmaier, Stefan Scheu, Leti Sundawati, Teja Tscharntke, Delphine Clara Zemp, Nathaly R. Guerrero-Ramírez

## Abstract

How to enhance biodiversity in monoculture-dominated landscapes is a key sustainability question that requires considering the spatial organization of ecological communities (beta diversity). Here, we experimentally tested if increasing landscape heterogeneity – through tree islands – is a suitable landscape restoration strategy when aiming to enhance multi-taxa diversity. We found that multi-taxa diversity resulted from islands fostering unique species (turnover: between 0.18 - 0.73) rather than species losses and gains (nestedness: between 0.03 - 0.34), suggesting that tree islands enhance diversity at the landscape scale. Through partial correlation networks, we revealed that landscape heterogeneity is associated with multi-taxa diversity (strength = 0.84). Soil biota were also central to the overall community by connecting beta diversity patterns across taxa. Our results show that increasing landscape heterogeneity enhances multi-taxa diversity in monoculture-dominant landscapes. Furthermore, we highlight that strategies aiming to enhance multi-taxa diversity should consider that spatial distributions of above- and below-ground communities are associated.

## Introduction

Habitat loss and degradation of natural ecosystems are major drivers of the global biodiversity crisis^1,2^, with more than half of the terrestrial land surface converted for anthropogenic uses^3^. Croplands have become the largest terrestrial land cover type on the planet^4^, with the net increase in tropical regions exceeding 100 million ha / decade^5^. Across the tropics, oil palm production has increased 15-fold in the last decades^6^, contributing significantly to land-use change and intensification and impacting global biodiversity hotspots. Specifically, oil palm plantations occupy 21 million hectares, mostly in Indonesia and Malaysia^7^. In the face of this biodiversity crisis, there is currently an unprecedented political will to restore degraded ecosystems and landscapes globally^8^. Therefore, it is fundamental to bring a complementary perspective to the United Nations (UN) on Ecosystem Restoration by expanding the restoration scope from degraded and abandoned lands to agricultural productive systems.

Embedding small patches of native trees (“tree islands”) in degraded landscapes is a promising strategy to enhance biodiversity and facilitate landscape restoration^9^. By actively planting trees or through natural regeneration, integrating natural habitats in monoculture-dominated landscapes can positively affect environmental heterogeneity^9–11^, where heterogeneous habitats can be associated with higher species diversity across taxa and spatial scales^12,13^. However, it remains uncertain to what extent environmental heterogeneity at the landscape-scale (i.e., landscape heterogeneity) can be leveraged to enhance the diversity of multiple taxonomic groups (i.e., multi-taxa diversity) in monoculture-dominated landscapes.

To inform conservation management and landscape restoration, it is essential to integrate insights from community assembly mechanisms; for example, through beta diversity that is the spatial distribution of ecological communities^14,15^. The assembly of ecological communities is determined by different factors, including biotic and abiotic filtering, environmental drift, and dispersal^16,17^. For instance, through direct and indirect species interactions, biotic filtering may play an important role in shaping biodiversity^18–20^ and the spatial organisation of (meta)communities^21–24^; explaining the growing interest in understanding the role of biotic interactions on community assembly in restoration contexts^25– 27^. Yet, our understanding of assembly mechanisms of multi-taxa communities in human-modified landscapes, particularly in the tropics, remains limited^15,28^.

Here, we assessed if multi-taxa diversity can be enhanced in large monoculture-dominated landscapes by embedding environmentally dissimilar tree islands. Furthermore, we investigated to what extent biotic associations are central to defining the spatial distribution of multi-taxa communities (i.e., multi-taxa beta diversity). To this end, we used comprehensive data from a unique tropical biodiversity enrichment experiment (EFForTS-BEE [Ecological and socio-economic functions of tropical lowland rainforest transformation systems: biodiversity enrichment experiment]^29^), located in Sumatra, Indonesia, a global hotspot of biodiversity loss^30^ and recent tropical deforestation^31^. Embedded within a 140-ha oil palm plantation, 52 experimental tree islands were planted with varying tree diversity and island size. In our study, we defined a landscape as “a geographical area, characterised by its content of observable, natural and human-induced, landscape elements” following^32^, with tree islands as the landscape elements (and no other surrounding land-use patches). This landscape-scale perspective with tree islands makes EFForTS-BEE unique among the largest network of tree diversity experiments worldwide (TreeDivNet^33^). We analysed multi-taxa diversity sampled three to five years after establishment, when the tree islands substantially differed in vegetation structural complexity^34^. We calculated beta diversity and its turnover and nestedness components (i.e., species losses and gains) using community data of understorey arthropods, soil biota (fungi, bacteria, and fauna), herbaceous plants, and trees (excluding planted trees). We expected that tree islands, varying in vegetation structural complexity (as a result of differences in island size and planted diversity^35^) and soil conditions, will increase total landscape diversity (i.e., gamma diversity) by fostering unique species resulting in higher turnover rather than species losses and gains, i.e., nestedness (Fig. 1).

**Figure 1.**
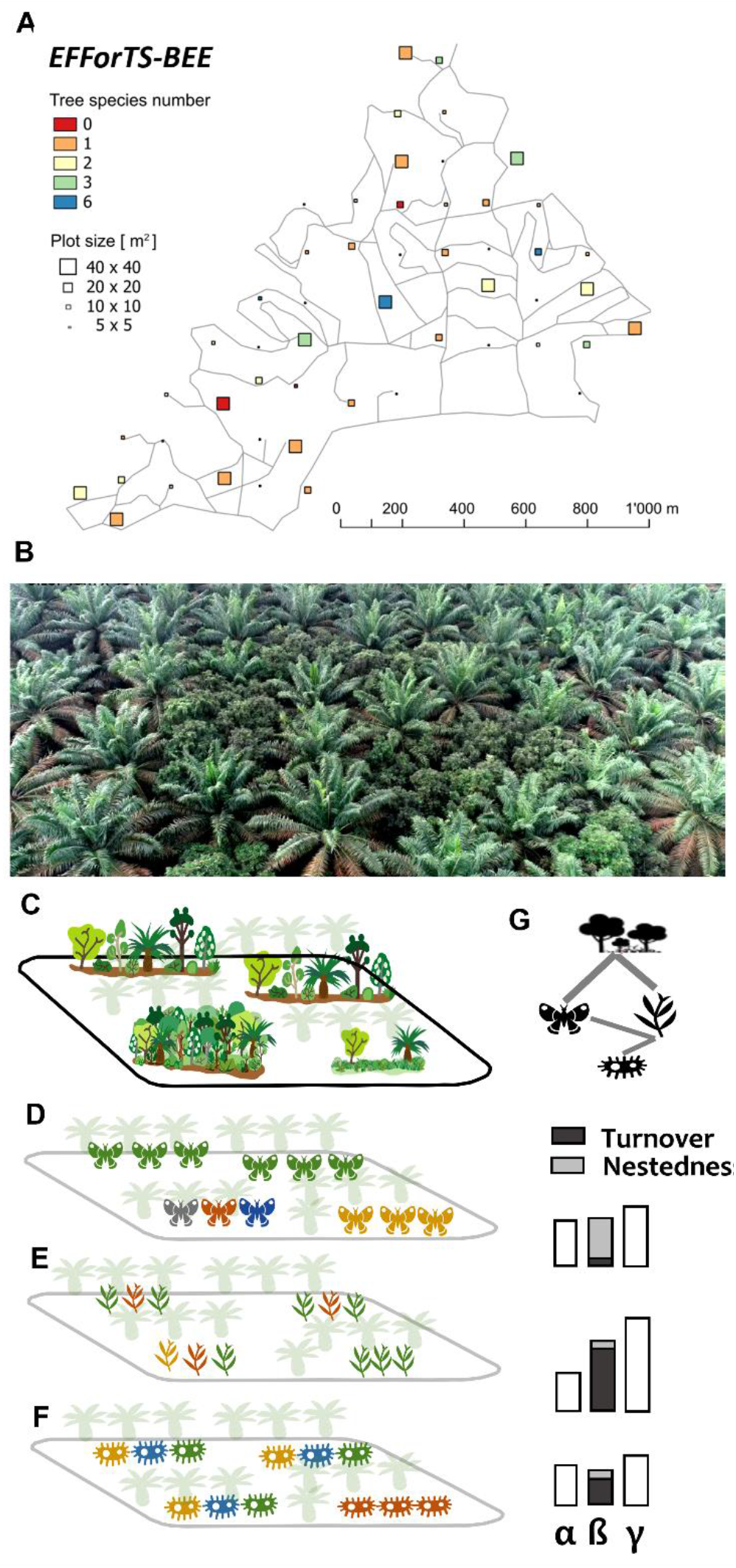
Tropical biodiversity enrichment experiment (EFForTS-BEE) and conceptual figures. **(A)** 52 experimental tree islands were established embedded within a 140-ha oil palm plantation, tree islands varying in tree native planted diversity and island size; **(B)** example of a tree island using a drone image; **(C)** if multi-taxa beta diversity is driven by habitat differentiation, higher landscape heterogeneity (resulting from islands differing in their vegetation structural complexity) is expected to be associated with beta diversity of multiple taxa. In contrast, if multi-taxa beta diversity is driven mostly by stochastic processes such as dispersion, landscape heterogeneity may not be associated with changes in beta diversity. Changes in beta diversity may be underlying by turnover, with higher turnover resulting in higher gamma diversity or, by nestedness (i.e., gain and species losses in light grey). Positive associations between landscape heterogeneity and beta diversity translate into greater dissimilarity in vegetation structural complexity between islands being associated with dissimilar multi-taxa communities e.g., **(C, D)** landscape heterogeneity and understory arthropods and **(C, E)** landscape heterogeneity and herbaceous plants. **(E-F)** A positive association between beta diversity of two taxa (e.g., herbaceous plants and soil bacteria) implies that tree islands that differ in herbaceous plant composition also differ in soil bacteria composition. **(G)** In the network, the nodes represent landscape heterogeneity and beta diversity (or one of its two components) for each taxon, and the links represent associations between the nodes.

To reveal the factors shaping the spatial distribution of multi-taxa communities (beta diversity, turnover and nestedness) across tree islands, we used partial correlation networks, which quantify associations among landscape heterogeneity (i.e., dissimilarity in vegetation structural complexity and soil conditions across tree islands) and beta diversities (or its underlying components) across taxa. Partial correlations can provide insights about associations shaping the spatial organisation of communities across taxa, e.g., similar niche requirements, dispersal limitations, and potential biotic interactions due to co-occurrences; with this approach particularly helpful in hyperdiverse regions such as the tropics, where biotic interactions likely structure strongly community assembly^21^ but assessing interactions is extremely challenging^36,37^. In the network, the nodes represent landscape heterogeneity and beta diversity (or one of its two components) for each taxon, and the links represent associations between the nodes. For example, positive associations between landscape heterogeneity and beta diversity translate into greater dissimilarity in vegetation structural complexity between islands being associated with dissimilar multi-taxa communities. A positive association between beta diversity of two taxa (e.g., herbaceous plants and soil bacteria) implies that tree islands that differ in herbaceous plant composition also differ in soil bacteria composition. Similarly, a positive association between turnover (or nestedness) between herbaceous plants or soil bacteria implies that tree islands that foster unique species (or are driven by species losses and gains) for herbaceous plants also show the same pattern(s) for soil bacteria (Fig 1).

## Results and discussion

### Gamma and beta diversity across tree islands embedded in an oil palm plantation

Across the 52 tree islands, we recorded 958 morphospecies of understorey arthropods, 8,159 operational taxonomic units (OTUs) of soil fungi, 47,856 OTUs of soil bacteria, 27 taxonomic groups of soil fauna (Supplementary Table S4), 75 herbaceous plant species, and 50 trees species - excluding planted trees (gamma diversity; all classifications are referred to as “species” below). Overall, across the 52 tree islands, beta diversity (calculated as Jaccard pairwise dissimilarity) varied among taxa, ranging from 0.31 for soil fauna to 0.77 for understorey arthropods. Beta diversity was mainly driven by species turnover, while nestedness, except for trees and soil fauna, played a minor role (Fig. 2). Specifically, the highest species turnover was found for soil fungi, understorey arthropods, and soil bacteria, accounting for ∼ 94% of the total beta diversity. Species turnover was lowest for trees (52%) and soil fauna (59%). We did not find major differences in the results when calculating beta diversity using Sørensen pairwise dissimilarity (Supplementary Figures S2 and S5). Hence, our results consistently indicate that beta diversity is primarily associated with the uniqueness of species assemblages rather than smaller assemblages being a subset of larger ones. Consequently, promoting the uniqueness of species assemblages with multiple tree islands appears as a promising strategy for enhancing biodiversity in monoculture-dominated landscapes, at least during the first years after tree island establishment.

**Figure 2.**
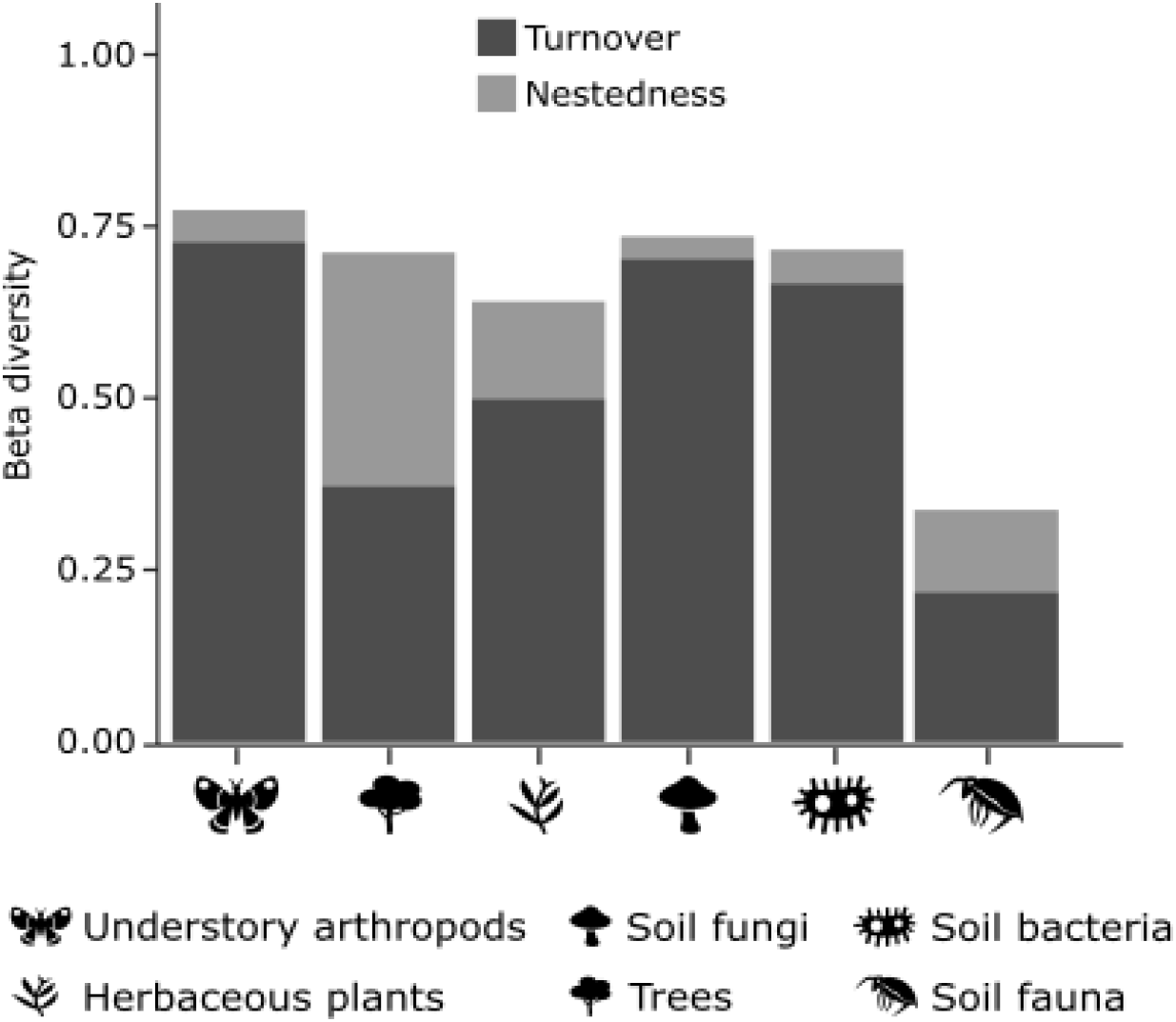
Turnover and nestedness components of beta diversity for taxonomic groups calculated with Jaccard index. Similar results were found when beta diversity was calculated using Sørensen pairwise dissimilarity (Supplementary Fig. S2).

The differences in beta diversity across taxa that our study revealed, might be explained by ecological processes related to dispersal ability, body size and life history. For instance, due to the long lifespan of trees, the influence of processes such as local extinction and colonisation may require more time than for other taxa. Furthermore, tree beta diversity patterns may be shaped mainly by seed sources in the surrounding landscape and by tree species with higher dispersal capacities^38^, explaining the unexpected high nestedness in human-modified ecosystems compared to tropical forests for trees^39^. While we expect overall patterns to hold, the influence of differences in sample coverage across taxa - particularly incomplete coverage for highly diverse taxon such as fungi - in terms of turnover and nestedness under- or over-estimations remains unknown. Finally, taxonomic resolution may impact our ecological understanding^40^, particularly for soil fauna that mainly was assessed at the level of orders (that often represent functional groups^41^). Contrasting resolutions reflect the challenge of biodiversity assessment in the species-rich tropics^36^. Despite that, soil fauna was a good indicator of overall multi-taxa community dissimilarity (see below). Therefore, we expect this crucial role to remain or be strengthened with higher resolution, but increases in resolution will likely result in higher beta diversity due to higher turnover.

### Insights of multi-taxa beta diversity through landscape heterogeneity and biotic associations

Beta diversity patterns across multiple taxa were correlated, with the network for beta diversity comprising 17 edges (Fig. 3A, Supplementary Table S6). The most connected taxa were soil fauna and bacteria (strength, i.e., the sum of absolute edge weights, = 0.82 and 0.71, with five and four edges with other nodes, respectively; Fig. 4A). By contrast, trees were the least connected (strength = 0.46, with four edges). The highest correlation coefficient was observed between soil fungi and bacteria beta diversity (+0.25). Turnover patterns for multi-taxa diversity were also correlated, with the network for turnover comprising eight edges (Fig. 3B, Supplementary Table S7). In the case of turnover, turnover of soil fauna and understorey arthropods were disconnected from the network. In other words, neither turnover patterns of soil fauna nor understory arthropods follow dis(similar) turnover patterns of other taxa, neither were associated with landscape heterogeneity. Finally, nestedness patterns for multi-taxa diversity were correlated except for trees (Fig. 3C, Supplementary Table S8), with the network retaining six edges. Yet, the nestedness network had low stability. Together, these results suggest that direct and indirect associations shape the spatial organisation of communities across taxa in tropical human-modified landscapes, supporting previous studies in temperate ecosystems^23,24^.

**Figure 3.**
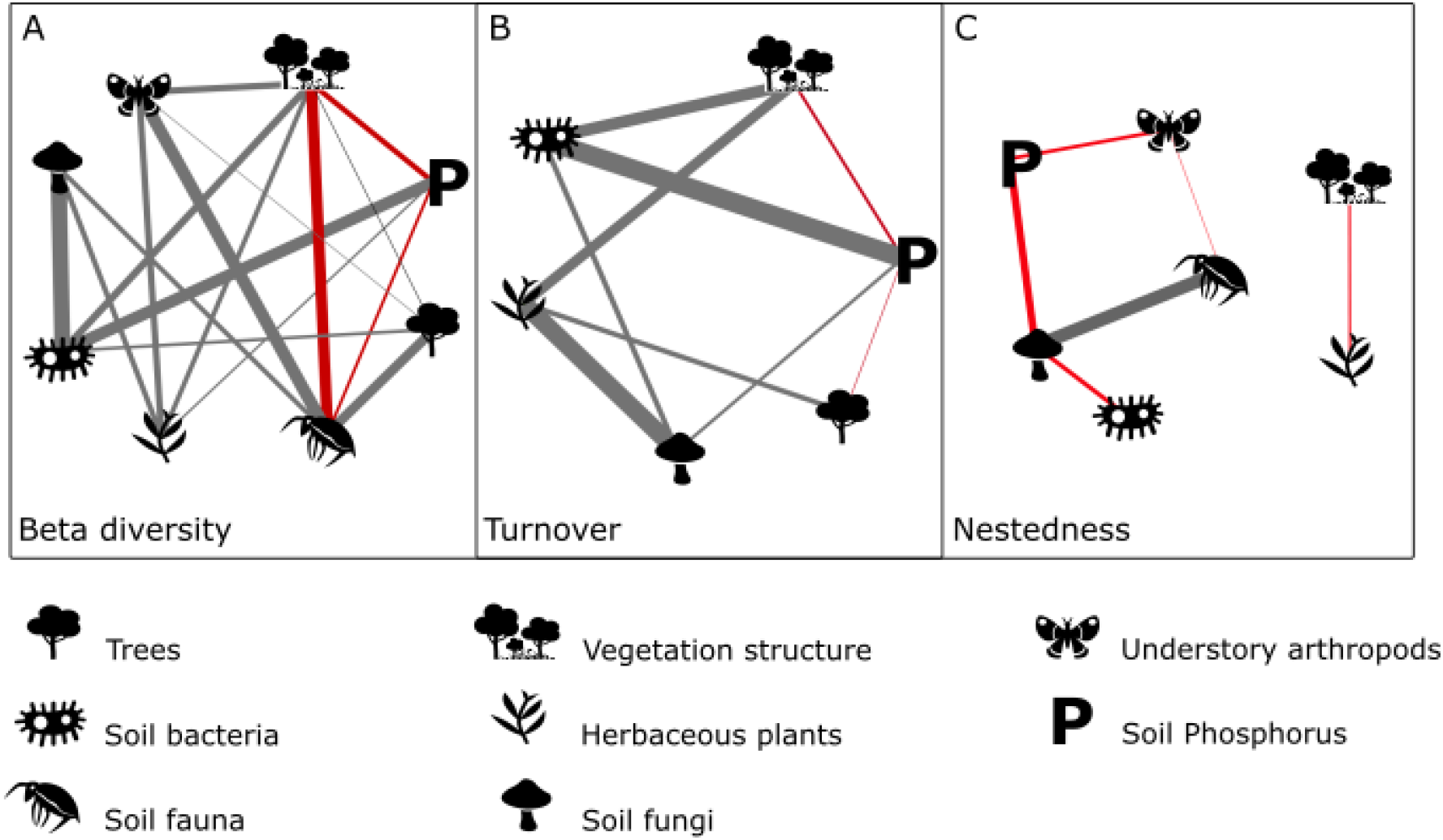
The role of landscape heterogeneity and biotic associations shaping multi-taxa beta diversity. Nodes represent **(A**) total beta diversity, **(B)** turnover, and **(C)** nestedness of multiple taxa and dissimilarity in vegetation structural complexity and soil phosphorus. Edges thicknesses, i.e., line thickness, are proportional to partial correlation coefficients, with grey and red edges representing positive (i.e., greater dissimilarity in vegetation structural complexity between islands being associated with dissimilar multi-taxa communities or tree islands that differ in composition for a taxon also differ in composition for another taxon) and negative (i.e., greater dissimilarity in vegetation structural complexity between islands being associated with similar multi-taxa communities or tree islands that differ in community compositions for a taxon have similar community compositions for another taxon) correlations, respectively. Edge length is not meaningful. Nodes with partial correlation coefficients equal to or near zero are not included in the corresponding network.

**Figure 4.**
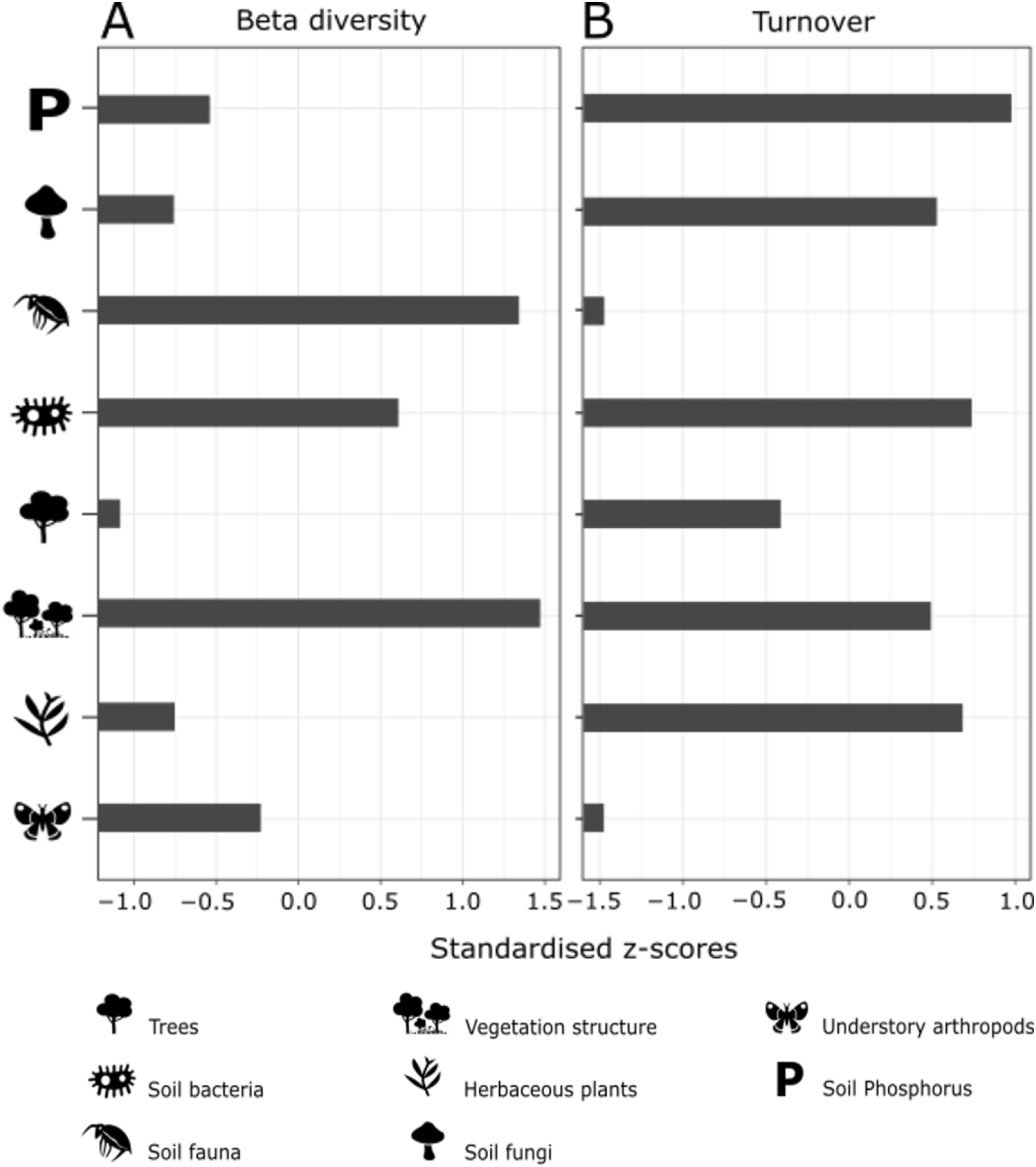
Importance of the individual taxa and landscape heterogeneity in shaping multi-taxa beta diversity. The centrality value (x-axis) for each node (y axis) is presented. Nodes represent **(A)** the total beta diversity and **(B)** turnover of multiple taxa and dissimilarity in vegetation structural complexity and soil phosphorus. The centrality value is quantified by the strength (i.e., the sum of absolute edge weights) in the undirected partial correlation networks and shown as standardised z-scores. Negative values indicate low centrality, whereas positive values indicate high centrality. Correlation stability coefficients of strength for beta diversity and turnover were 0.36 and 0.44, respectively. For nestedness, the correlation stability coefficient was lower than 0.25, suggesting lower stability of this network that is therefore not presented in this figure (see Supplementary Fig. S3). Other centrality measures, i.e., betweenness and closeness, are shown in Supplementary Fig. S3. Observed and non-parametric bootstrap mean and 95% CI estimated are shown in Supplementary Fig. S8 and S9.

Our results point toward the key role of below-ground organisms in structuring multi-taxa beta diversity patterns. Soil biota (soil fauna, bacteria, and fungi) are central to the overall ecological community as its beta diversity patterns are associated with beta diversity patterns of other taxonomic groups and with abiotic variables (for different centrality indices, Supplementary Fig. S3). Soil biota may act as an indicator of current conditions, the result of legacy effects from previous land-uses (e.g., oil palm plantation or tropical forest), or both^42^. For example, soil fauna composition can be associated with differences in specific organic materials (reflecting the heterogeneity before the land-use conversion) and time delays because of the limited dispersion of soil fauna^43^. Similar beta diversity patterns between soil fauna and soil fungi may be underlain by species interactions (e.g., soil fungi as an important food source in soil food webs^44^), similar niche requirements and/or dispersal limitations influencing soil biota (symbiotroph, pathotroph and saprotroph, Supplementary Fig. S4 – S7; Supplementary Tables S9 – S11). Associations between soil biota and trees can result from plant-soil feedbacks, with soil fauna potentially influencing vegetation dynamics and above-ground biodiversity^45^. For instance, soil biota have been shown to affect understorey arthropods (particularly pollinators, Supplementary Fig. S4 – S6) when soil biota indirectly affect floral traits (e.g., bacteria, root herbivores, and mycorrhizal fungi), influencing pollination attractions and plant fitness^46^. While detailed plant-soil feedback experiments would be required to disentangle the mechanisms of above- and below-ground associations shaping multi-taxa dynamics, here we provide further evidence highlighting the importance of integrating the belowground compartment towards elucidating dynamics in monoculture-dominated landscapes.

Landscape heterogeneity played a crucial role in all three networks (Fig. 3). For instance, dissimilarity in vegetation structural complexity was the most connected node (strength = 0.84 with four edges to other nodes) in the beta diversity network. Besides, soil P was the most connected node (strength = 0.49 with four edges, Fig 4B) in the species turnover network. The highest and lowest correlation of soil P was found with soil bacteria and fungi beta diversity, respectively (+0.18 and +0.11). This suggests that landscape heterogeneity can promote beta diversity by fostering different species compositions, reinforcing the role of enriched tree islands in influencing community assemblages and biodiversity at the landscape-scale (i.e., beta and gamma diversity). Further, it implies that dissimilarity in abiotic conditions can directly or indirectly impact multiple taxa. The influence of vegetation structural complexity on multi-taxa diversity may act *via* altering light and microclimatic conditions^47^ and other characteristics associated with variation in local planted tree species diversity and identity – with both shaping vegetation structural complexity^35^. Furthermore, the influence of tree islands on multi-taxa diversity might reflect the removal of environmental filtering associated with conventional management such as liming and fertilisation, which is responsible for biotic homogeneity in monoculture-dominated landscapes. Further possible mechanisms include enhanced nutrient cycling and plant litter decomposition^48,49^, particularly in ecosystems under transition (e.g., primary or secondary succession)^50^.

### Conclusions

We conclude that enriching monocultures with tree islands varying in vegetation structural complexity (as a result, for instance, of tree planting diversity and island size) can foster unique ecological communities above- and below-ground and thereby promote multi-taxa diversity at the landscape-scale (beta and gamma diversity). Additionally, we suggest distributing tree islands across the monoculture-dominated landscape to enhance multi-taxa diversity by capturing contrasting soil conditions. Landscape restoration strategies aiming to enhance multi-taxa diversity should consider not only key abiotic conditions but also the extent to which biotic associations play an important role in shaping ecological communities at landscape-scale. By enhancing biodiversity at the landscape level in monoculture-dominated tropical landscapes, we bring a complementary perspective to the UN Decade on Ecosystem Restoration and provide experimental evidence urgently needed to guide interventions for landscape restoration in productive agricultural systems.

## Materials and Methods

### Study area

This study was conducted in the Biodiversity Enrichment Experiment (EFForTS-BEE) located in Jambi province, Sumatra, Indonesia. The main aim of EFForTS-BEE is to evaluate the potential of establishing tree islands^9^ within an industrial oil palm plantation as a restoration measure to enhance biodiversity and ecosystem functioning while maintaining financial benefits (^29^, Zemp et al., in revision). The area is characterised by a humid tropical climate with two peak rainy seasons (March and December) and a dryer period extending from July to August^29^. The mean temperature is 26.7 ± 1.0 °C, and the mean annual precipitation is 2235 ± 385 mm (1991 - 2011). The predominant soil type in the region is loamy Acrisol^51^. EFForTS-BEE was established in December 2013 and consists of 52 experimental plots, i.e., tree islands varying in plot size of 25 m^2^, 100 m^2^, 400 m^2^, and 1,600 m^2^, and planted tree diversity level 0, equal to no tree planted, 1, 2, 3, and 6 tree species planted in a plot, all embedded in a 140-ha oil palm plantation (01.95° S and 103.25° E, 47 ± 11 m a.s.l.). The experiment follows a random partition design aiming to disentangle the linear effects of tree diversity and plot size and the non-linear effects of tree species composition^29^. For details of the experimental design, see ref^29^. The planted species represent native, multi-purpose trees used for the production of fruits (*Parkia speciosa* Hassk, *Archidendron* jiringa (Jack) I.C.Nielsen, and *Durio zibethinus* L.), timber (*Peronema canescens* Jack, and *Shorea leprosula* Miq.), and natural latex (*Dyera polyphylla* (Miq.) Steenis)^34^.

### Data collection

The data for this study were collected between October 2016 and May 2018. We sampled above-ground and below-ground taxa, including understorey arthropods, soil biota (soil fungi, soil bacteria, and soil fauna), herbaceous plants, trees, vegetation structural complexity measures, and soil conditions, with all measurements within the 52 tree islands, i.e., plots. Arthropods sampled at the height of the understorey vegetation (referred to as “understorey arthropods”) were sampled three times with six pan traps (2 × 3 pan traps) equally distributed per plot, for 45 hours from October 2016 to January 2017. The traps were made of white plastic bowls coloured with yellow UV paint^52^ and filled with water and a drop of detergent. All individuals were preserved in 70% Ethanol, sorted by morphospecies, and subsequently identified into higher taxonomic classification possible (i.e., 14 groups/families) and their corresponding functional groups (e.g., Table S5).

Soil biota and herbaceous plants were surveyed in the same subplot of 5 × 5 m area established within each plot^29^. Specifically, soil fungi were sampled and collected in December 2016 from three soil cores per plot (10 cm depth and 4 cm diameter) and identified through DNA extraction and next-generation sequencing^42^. OTUs were classified taxonomically using the *BLAST* algorithm (blastn, v2.7.1; ^53^) and the UNITE v7.2 (UNITE_public_01.12.2017.fasta; ^54^). Soil bacteria were obtained for each subplot from three 10 cm cores of topsoil, placed at 1 m far from the adjacent trees. The soil cores were mixed, homogenised and cleared from roots before DNA and RNA extraction and posterior classification^55^. In each plot, soil fauna communities were assessed in four soil samples of 16 × 16 cm using a spade down to a depth of 5 cm plus the entire overlying litter layer. The animals extracted from the soil samples by heat were counted and classified into taxonomic groups, corresponding to key functional soil invertebrate guilds (mainly groups/families, Supplementary Table S4)^41,56,57^. Herbaceous plants, described as all non-woody plants lower than 1.3 metres in height, were identified from February to March 2018. Trees refer to all free-standing woody plants with a minimum height of 1.3 m, inventoried in the total area of the experimental tree islands in August 2018, excluding the trees planted at the onset of the experiment.

Soil nutrient variables, including total carbon (C) and nitrogen (N) concentration (g mg^-1^), C-to-N ratio, and plant-available P concentration (mg g^-1^), were quantified using the same soil samples as for soil fungi collected in December 2016 (see below). Total C and N were determined via the combustion method in a C/N analyser^42^. Plant-available P was quantified following Bray & Kurtz^58^. The soil samples were mixed with Bray-I Extraction solution, shaken for 60 min, and filtered with phosphate-free filters. P concentration of filtrates was measured using inductively coupled plasma mass spectrometry^42^.

We quantified vegetation structural complexity using multiple terrestrial laser scans taken between September and October 2016^35^. We calculated Effective Number of Layer (ENL), which describes the vertical structure of forest stands and is influenced by the stand height and the vegetation distribution across vertical layers^59^. In addition, we calculated Mean Fractal Dimension (MeanFRAC) and Stand Structural Complexity Index (SSCI). MeanFRAC is defined as the arithmetic mean of fractal dimensions describing the geometric complexity of the stand^60^. MeanFRAC is associated with enriched tree island conditions, i.e., planted tree composition, richness, and tree island size^35^. SSCI describes the arrangement of tree components in three-dimensional space^60,61^.

### Beta diversity and landscape heterogeneity

For each taxa, beta diversity was calculated using species incidence-based pairwise dissimilarity matrices (presence-absence data) with the function *beta*.*pair* from the package *betapart* version 1.5.4^62^. We partitioned beta diversity into turnover and nestedness components^62,63^. The Jaccard pairwise dissimilarity (β_jacc_) among plots was computed as β_jacc_ = β_jtu_ + β_jne_, where β_jtu_ accounted for the turnover fraction of Jaccard pairwise dissimilarity, and β_jne_ accounted for the nestedness-resultant dissimilarity fraction. We calculated beta diversity using community data (incl. operational taxonomic units, taxonomic groups, morphospecies or species – referred as species in the text). In addition, we calculated beta diversity using Sørensen pairwise dissimilarity, which incorporates turnover and richness differences as β_sor_ = β_sim_ + β_sne_. In this case, β_sim_ accounted for turnover measured as Simpson pairwise dissimilarity, and β_sne_ accounted for the patterns of beta diversity causing nestedness, measured as the nestedness-resultant dissimilarity fraction of Sørensen dissimilarity (Supplementary Fig. S2 and S5). While Jaccard considers the proportion of unique species in the entire pool, Sørensen considers the proportion of unique species per site^64^.

For the abiotic variables (vegetation structural complexity and soil conditions), we calculated pairwise dissimilarity between all matrix rows, i.e., tree islands, using the function *dist* from the R stats package. We used the Euclidean distance method, calculated as a true straight-line distance between all matrix rows in Euclidean space.

Multivariate normality was tested with Mardia’s multivariate skewness and kurtosis coefficients using the function *mvn* from the R package *MVN* version 5.9^65^. When the test did not state multivariate normality, a non-paranormal transformation to achieve Gaussian distribution was implemented using the function *huge*.*npn* and the setting *shrinkage* based on a shrunken Empirical Cumulative Distribution Function (ECDF) from the R package *huge* version 1.3.5^66^.

### Partial correlation networks

We applied partial correlation networks to study associations between landscape heterogeneity and the beta diversity (turnover or nestedness) among multiple taxa. An association between taxa indicates the covariation of the spatial distribution of ecological communities among taxa. Advantages of partial correlation networks are threefold: first, they describe correlations between a set of conditionally independent variables^67^, second, they do not require *a priori* knowledge of the structure^68^; and finally, the correlations can be graphically represented and analysed to reveal key interdependencies and highly connected variables^69^. Partial correlation networks have been widely used to infer pairwise species interactions from observed presence-absence matrices^68^. A network is composed of nodes and edges, where the nodes represent the beta diversity (or turnover or nestedness) of the different taxa and the dissimilarity of vegetation structural complexity and soil conditions. The edges (i.e., links connecting pairs of nodes) represent correlations between nodes, in our case, undirected partial correlation coefficients^23^. Edges can be either positive or negative correlations (representing the covariation of the spatial distribution of ecological communities between taxa), and can be absent, indicating no or weak correlation between a set of variables^70^. When positive, the (dis)similarity in species composition between tree islands changes in the same direction for both taxa, when negative, the (dis)similarity in species composition for a taxon increases while it decreases for the other taxon.

We used the graphical lasso method (Least Absolute Shrinkage and Selection Operator) as implemented in the R package *bootnet* version 1.4.3^71^ to build and analyse the networks. This method displays the unconditional association between two nodes once the influence of other variables is controlled (i.e., partial correlations^67^), reducing the risk of spurious relationships that can emerge from multicollinearity^70^. The Lasso method applies a regularisation penalty using a tuning parameter to reduce the number of parameters displayed. As a result, only a small number of partial correlations (i.e., the highest values) are used to explain the interconnections among variables^67^. We selected the tuning parameter with the Extended Bayesian Information Criterion EBIC^72^ using the function *EBICglasso* from the package *qgraph* version 1.6.9^73^ (tuning parameter = 0.5). The partial correlations were represented graphically in networks with undirected weighted edges (i.e., there is an association, but the direction is not determined) using *ggraph* R package version 2.0.5^74^. With the weighted networks, we consider the correlations among nodes and the weight of these correlations (partial correlation coefficients^75^).

We tested the influence of different abiotic variables on network connectivity. To do so, we included various combinations of vegetation structural complexity metrics and soil conditions and measured the resulting number of edges in the network and the proportional changes. We found the highest network connectivity when MeanFRAC and soil P were included (Supplementary Tables S2 and S3). Other structural metrics or soil conditions did not increase network connectivity and were highly correlated with other environmental variables (Supplementary Table S1 and Fig. S1). Therefore, we only included MeanFRAC (named hereafter as vegetation structural complexity) and soil P in the final networks presented in this study.

We quantified the importance of specific nodes (i.e., certain taxon or a particular environmental variable) for structuring or maintaining the overall (i.e., multi taxa) network by calculating three centrality measures commonly used in complex network approaches strength, betweenness, and closeness. Strength is the sum of absolute edge weights that a node has with the others^67^. The higher the strength value of a node, the higher the influence it has on influencing the composition and structure of the community^24^. Betweenness looks at the proportion of shortest paths between any pair of nodes that pass through a specific node. The shortest path is defined as the path with the minimum distance (calculated by adding the edges’ weights) needed to connect two nodes. Hence, a node with high betweenness lies “in-between” other nodes’ shortest paths in the network. High betweenness indicates that a node plays a crucial role in the connectivity and stability of the network, for example, implying a cascading effect with large consequences on the overall network when the node is lost^76^. Closeness describes the undirected connectance of a node to the other nodes in a network, calculated as the average distance of the shortest path from a specific node to all other nodes^67^. Because of its proximity to all other nodes, the node with the highest closeness centrality plays a crucial role in the overall network^76^ (Supplementary Fig. S3 and S7).

The accuracy of the parameters and measures estimated in a network depends greatly on sample size and variability^75^. Thus, we assessed the accuracy of the different networks (i.e., sensitivity to sampling variation) by estimating confidence intervals on the weight of the edges with a non-parametric bootstrapping of 1000 samples, with a confidence interval of 95%^75^, using the *bootnet* R package version 1.4.3^71^. To assess the stability of centrality indices, we used a case-dropping subset bootstrap from the package *bootnet*^71^. We calculated the correlation stability coefficient (CS-coefficient), which represents the maximum number of observations that can be dropped (in at least 95 % of the samples) so that the correlation between original centrality indices and the indices re-calculated with a subset of the data is 0.7 or higher^67^. The threshold considered stable for the CS-coefficient should be no less than 0.25 and desirable higher than 0.5. Results of the sensitivity analysis are presented in Supplementary Fig. S8-S13.

Data were analysed with the software environment R, version 4.1.1 (R Development Core Team, 2021), using the packages *ade4*^77^, *betapart*^62^, *bootnet*^71^, *data*.*table*^78^, *ggplot2*^79^, *ggraph*^74^, *glasso*^80^, *huge*^66^, *igraph*^81^, *MVN*^65^, *plyr*^82^, *qgraph*^73^, *reshape2*^83^, *rlist*^84^, *tidyverse*^85^, and *vegan*^86^. Our code is based on the R code provided by Ohlmann e*t al*. (2018)^23^.

## Data availability

The data and code to reproduce the results will be available on Zenodo.

## Acknowledgements

We thank all scientists who contributed to the data analysed in this study, including Prof. Dr. Mark Maraun. We also thank PT Humusindo Makmur Sejati and Pak Hasbi and his family for granting us access to and use of their properties. We thank the many field assistants, in particular, Juliandi and Eduard J. Siahaan, for their support in the field. We are grateful for the logistical support by the EFForTS staff and coordination. This study was financed by the Deutsche Forschungsgemeinschaft (DFG) German Research Foundation – project number 192626868 – SFB 990 in the framework of the collaborative German – Indonesian research project CRC990 EFForTS (http://www.unigoettingen.de/crc990). Research permit by the Indonesia Ministry of Research and Technology (337/SIP/FRP/E5/Dit.KI/IX/2016). Nathaly R. Guerrero-Ramírez thanks the Dorothea Schlözer Postdoctoral Programme of the Georg-August-Universität for their support. EFForTS-BEE is a member of the global network of tree diversity experiments TreeDivNet (http://www.treedivnet.ugent.be).

## Author Contributions

V.M-S., H.K., D.H., D.C.Z., and N.R.G-R. designed research; I.A., J.B., D.B., F.B., A.K., A.P., and L.S., collected data with supervision from H.K., R.D., I.G., D.H., A.P., S.S, T.T, D.C.Z; V.M-S. analysed data with assistance from D.C.Z. and N.R.G-R.; and V.M-S., D.C.Z., and N.R.G-R. wrote the paper with assistance from H.K., I.A., J.B., D.B., F.B., R.D., I.G., J.H., D.H., B.I., A.K., A.P., A.P., L.S., S.S., L.S., T.T.

## Competing Interests

The authors declare no competing interest.

## Materials and correspondence

Correspondence and materials requests should be addressed to Vannesa Montoya-Sánchez.

## Notes

### Competing Interest Statement

The authors have declared no competing interest.

