## Supplementary information for "Landscape heterogeneity and soil biota are central to multi-taxa diversity for landscape restoration"

### Supplementary Information Text

### Supplementary methods

Understorey arthropods and soil fungi were classified into functional groups. For understorey arthropods, Hymenoptera, Lepidoptera and Araneae were classified into pollinator, parasitic, or predatory understorey arthropods (representing 64% of total understorey arthropods data), using different identification keys[1–9]. Soil fungi were classified into pathotroph, saprotroph and symbiotroph using the FunGuild database based on taxonomic affiliations (representing 31% of total soil fungi data; [10]).

### Supplementary results

**Beta diversity for functional groups across tree islands embedded in an oil-palm dominated landscape:** For understorey arthropods classified into functional groups, turnover was 98% in parasitic and predators, and 87% in pollinators. For soil fungi, turnover was 95% in pathotroph and symbiotroph and 93% in saprotroph (Fig S4 and S5).

**Insights of multi-taxa beta diversity of functional groups through landscape heterogeneity and biotic associations:** Beta diversity patterns of different functional groups are not independent of other functional and/or taxonomic groups (except parasitic understorey arthropods). Relationships among functional groups were mostly observed for functional groups of fungi (Fig S6A), with the strongest partial correlation (i.e., the highest edge value) in the network between saprotroph fungi and symbiotroph fungi (+0.45). Further, saprotroph fungi had the highest strength (0.78), followed by soil fauna (0.76) and symbiotroph fungi (0.74). Centrality stability (CS) analysis indicates that node strength (CS (cor = 0.7) = 0.52) is stable. Closeness and betweenness centralities were less stable and should be interpreted with care (CS (cor = 0.7) = 0 and CS (cor = 0.7) = 0.05, respectively; Fig S7A).

For turnover, the strongest partial correlation was found between pathotroph and saprotroph fungi (+0.31), followed by saprotroph fungi and symbiotroph fungi (+0.21) (Fig S6B). The three nodes with the highest strength were saprotroph fungi (0.51), pathotroph fungi (0.49), and symbiotroph fungi (0.39), forming a three-node network with no connection with the rest of the taxonomic groups or environmental distances. Negative partial correlations were observed between the turnover of trees and pollinators (-0.12), trees and parasitic understorey arthropods (-0.11), and herbaceous plants and pollinators (-0.11). According to the centrality stability analysis, centrality metrics should be interpreted with caution as the stability values calculated are below the limits to be considered stable (betweenness CS (cor = 0.7) = 0, closeness CS (cor = 0.7) = 0, and strength CS (cor = 0.7) = 0.28; Fig S7B).

Relationships between functional groups of fungi shaped the network for nestedness, in which only three partial correlations were observed (Fig S6C). Pathotroph fungi connected to saprotroph fungi (+0.37) and symbiotroph fungi (+0.17). Centrality measurements revealed pathotroph fungi as the node with the highest strength (0.54) (Fig S7C). Saprotroph fungi (0.37) and symbiotroph fungi (0.17) were the nodes with the second and third strongest strength, respectively. Centrality stability analysis indicates that node strength (CS (cor = 0.7) = 0.75) is stable. Closeness and betweenness centralities were less stable and should be interpreted with care ((CS (cor = 0) and (CS (cor = 0.44), respectively; Fig S7C).

### Supplementary tables

**Supplementary Table S1.** Correlations coefficients and p-values between vegetation structural complexity measures and soil nutrient conditions (using Pearson correlation). P-values are in grey, significant correlations coefficients (p-values <0.05) in bold. MeanFRAC: Mean fractal dimension; SSC: Stand Structural Complexity Index; ENL: Effective Number of Layers.

| | Soil C | soilN | soilP | soilCN | SSC | ENL | MeanFRAC |
|---|---|---|---|---|---|---|---|
| **soilC** | 1 | 0.00 | 0.30 | 0.82 | 0.31 | 0.16 | 0.03 |
| **soilN** | **0.95** | 1 | 0.30 | 0.01 | 0.21 | 0.10 | 0.14 |
| **soilP** | -0.15 | -0.15 | 1 | 0.67 | 0.29 | 0.26 | 1 |
| **soilCN** | -0.03 | **-0.35** | 0.06 | 1 | 0.47 | 0.30 | 0.24 |
| **SSC** | 0.14 | 0.18 | 0.15 | -0.10 | 1 | 0.75 | 0.00 |
| **ENL** | -0.20 | -0.23 | 0.16 | 0.15 | 0.04 | 1 | 0.00 |
| **MeanFRAC** | **0.30** | 0.34 | 0.00 | -0.17 | **0.77** | **-0.57** | 1 |

**Supplementary Table S2.** Variation in connectivity of the network for beta diversity, turnover and nestedness by including contrasting metrics of vegetation structural complexity and soil conditions. Connectivity of the network (in relative value) is calculated as the total number of observed partial correlations (positive and negative) in the network (inferred using the Glasso algorithm) divided by the total number of possible correlations calculated as Pearson correlations (inferred as the inverse of the empirical variance-covariance matrix).

| | ***Connectivity of the network (relative value)** | | |
|---|---|---|---|
| **Variables** | **Beta diversity** | **Turnover** | **Nestedness** |
| soil nitrogen (N) – Mean fractal dimension (MeanFRAC) | 0.500 | 0.250 | 0.143 |
| soil carbon (C) - MeanFRAC | 0.536 | 0.250 | 0.143 |
| C-to-N ratio - MeanFRAC | 0.464 | 0.178 | 0.107 |
| soil phosphorus (P) - MeanFRAC | 0.607 | 0.321 | 0.178 |
| soil N – Stand structural complexity index (SSC) | 0.428 | 0.321 | 0.143 |
| soil C -SSC | 0.428 | 0.321 | 0.143 |
| C-to-N ratio – SSC | 0.392 | 0.250 | 0.142 |
| soil P – SSC | 0.392 | 0.321 | 0.178 |
| soil N – Effective number of layers (ENL) | 0.357 | 0.179 | 0.071 |
| soil C – ENL | 0.357 | 0.179 | 0.071 |
| C-to-N ratio - ENL | 0.393 | 0.107 | 0.071 |
| soil P - ENL | 0.357 | 0.214 | 0.142 |
| C-to-N ratio - soil P - MeanFRAC | 0.472 | 0.250 | 0.111 |
| C-to-N ratio - soil P - SSC | 0.333 | 0.277 | 0.166 |

| | *Connectivity of the network (relative value) | | |
|---|---|---|---|
| Variables | Beta diversity | Turnover | Nestedness |
| C-to-N ratio - soil P - ENL | 0.277 | 0.166 | 0.111 |

**Supplementary Table S3.** Number of undirected edges of the network for beta diversity, turnover and nestedness at different combinations of variables.

| | Number of undirected edges | | |
|---|---|---|---|
| **Variables** | **Beta diversity** | **Turnover** | **Nestedness** |
| Soil nitrogen (N) - Mean fractal dimension (MeanFRAC) | 14 | 7 | 4 |
| soil carbon (C) - MeanFRAC | 15 | 7 | 4 |
| C-to-N ratio - MeanFRAC | 13 | 5 | 3 |
| soil phosphorus (P) - MeanFRAC | 17 | 9 | 5 |
| soil N - Stand structure complexity index (SSC) | 12 | 9 | 4 |
| soil C -SSC | 12 | 9 | 4 |
| C/N ratio - SSC | 11 | 7 | 4 |
| soil P - SSC | 11 | 9 | 5 |
| soil N - Effective number of layers (ENL) | 10 | 5 | 2 |
| soil C - ENL | 10 | 5 | 2 |
| C-to-N ratio - ENL | 11 | 3 | 2 |

| | Number of undirected edges | | |
|---|---|---|---|
| **Variables** | **Beta diversity** | **Turnover** | **Nestedness** |
| soil P - ENL | 10 | 6 | 4 |
| C-to-N ratio - soil P - MeanFRAC | 17 | 9 | 4 |
| C-to-N ratio - soil P - SSC | 12 | 10 | 6 |
| C-to-N ratio - soil P - ENL | 10 | 6 | 4 |

**Supplementary Table S4.** Soil fauna and arthropod orders/families found in the 52 enriched tree islands

| | | | | | |
|---|---|---|---|---|---|
| 1 | Araneae | 11 | Gastropoda | 21 | Protura |
| 2 | Blattodea | 12 | Heteroptera | 22 | Pseudoscorpionida |
| 3 | Chilopoda | 13 | Hymenoptera | 23 | Psocoptera |
| 4 | Coleoptera | 14 | Isopoda | 24 | Schizomida |
| 5 | Collembola | 15 | Lepidoptera | 25 | Sternorrhyncha |
| 6 | Dermaptera | 16 | Lumbricina | 26 | Symphyla |
| 7 | Diplopoda | 17 | Mesostigmata | 27 | Thysanoptera |
| 8 | Diplura: Campodeidae | 18 | Opiliones | | |
| 9 | Diplura: Japygidae | 19 | Oribatida | | |
| 10 | Diptera | 20 | Orthoptera | | |

**Supplementary Table S5.** Groups/families found in the 52 enriched tree islands included in understory arthropods. Pollinator, parasitic, and predatory understorey arthropods were used for the analyses at functional group level

| **Group** | | **Guild** |
|---|---|---|
| Acarina | - Acariformes | Decomposers |
| Araneae | - Amaurobiidae, Araneidae, Lycosidae, Oxyiopidae, Salticidae, Thomisidae, Uloboridae | Predators |
| Araneae | - Others | Predators |
| Blattodea | | Decomposers |
| Coleoptera | - Carabidae, Coccinellidae, Scarabaedidae, Staphylinidae | Predators |
| Coleoptera | - Attelabidae, Chrysomelidae, Curculionidae, Mordellidae | Herbivores |
| Coleoptera | - Nitidulidae, Scolytidae | Decomposers |
| Coleoptera | - Others | |
| Collembola | | Decomposers |
| Dermaptera | | Decomposers |
| Diptera | - Asilidae | Predators |
| Diptera | - Stratiomyidae | Decomposers |
| Diptera | - Others | |
| Gastropoda | | |
| Hemiptera | - Alydidae, Aphididae, Cercopidae, Cicadellidae, Cydinidae, Delphacidae, Derbiidae, Flatidae, Fulgoridae, Lygaeidae, Membracidae, Pentatomidae, Reduviidae, Rhopalidae, Scutelleridae, Tingidae | Herbivores |
| Hemiptera | - Others | |
| Hymenoptera | - Aphelinidae, Braconidae, Ceraphronidae, Chalcididae, Diapriidae, Dryinidae, Encyrtidae, Eucoilidae, Eulophidae, Evaniidae, Figitidae, Ichneumonidae, Mymaridae, Platygastridae, Pteromalidae, Scelionidae, Tenthredinidae, Trichogrammatidae (*Trichogramma*) | Parasitica |
| Hymenoptera | - Apidae (*Amegilla zonata*), Colletidae, Halictidae (*Lasioglossum, Nomia* and Others) | Pollinators |
| Hymenoptera | - Bethylidae, Crabronidae, Formicidae *(Anoplolepis, Anaplolepis gracilipes, Crematogaster, Monomorium, Oecophylla*, Others *Paratrechina, Pheidole, Polyrhachis, Tapinoma, Technomyrmex*), Mutillidae, Pompilidae, Scoliidae, Sphecidae, Tiphiidae, Vespidae | Predators |
| Hymenoptera | - Others | |
| Lepidoptera | - Hesperiidae | Pollinators |

| Group | | Guild |
|---|---|---|
| Lepidoptera | - Others | |
| Orthoptera | - Acrididae, Alydidae, Gryllidae, Pyrgomorphidae, Tettigoniidae | Herbivores |
| Orthoptera | - Others | |
| Psocoptera | | Decomposers |
| Thysanoptera | | Herbivores |

**Supplementary Table S6.** Partial correlations coefficients for beta diversity of the Jaccard dissimilarity index for the taxonomic groups and the environmental distances calculated using EBIC graphical lasso.

| | SP | VS | UA | SFU | SB | HP | SFA | TR |
|---|---|---|---|---|---|---|---|---|
| Soil phosphorus (SP) | 0 | | | | | | | |
| Vegetation structure (VS) | -0.13 | 0 | | | | | | |
| Understorey arthropods (UA) | 0 | 0.14 | 0 | | | | | |
| Soil fungi (SFU) | 0 | 0 | 0 | 0 | | | | |
| Soil bacteria (SB) | 0.20 | 0.15 | 0 | 0.25 | 0 | | | |
| Herbaceous plant (HP) | 0.10 | 0.13 | 0.15 | 0.13 | 0 | 0 | | |
| Soil fauna (SFA) | -0.11 | -0.20 | 0.21 | 0.13 | 0 | 0 | 0 | |
| Trees (TR) | 0 | 0.10 | 0.10 | 0 | 0.11 | 0 | 0.18 | 0 |

**Supplementary Table S7.** Partial correlations coefficients for the turnover fraction of beta diversity of the Jaccard dissimilarity index for the taxonomic groups and the environmental distances calculated using EBIC graphical lasso

| | **SP** | **VS** | **UA** | **SFU** | **SB** | **HP** | **SFA** | **TR** |
|---|---|---|---|---|---|---|---|---|
| Soil phosphorus (SP) | 0 | | | | | | | |
| Vegetation structure (VS) | -0.11 | 0 | | | | | | |
| Understorey arthropods (UA) | 0 | 0 | 0 | | | | | |
| Soil fungi (SFU) | 0.11 | 0 | 0 | 0 | | | | |
| Soil bacteria (SB) | 0.18 | 0.15 | 0 | 0.12 | 0 | | | |
| Herbaceous plant (HP) | 0 | 0.14 | 0 | 0.18 | 0 | 0 | | |
| Soil fauna (SFA) | 0 | 0 | 0 | 0 | 0 | 0 | 0 | |
| Trees (TR) | -0.10 | 0 | 0 | 0 | 0 | 0.12 | 0 | 0 |

**Supplementary Table S8.** Partial correlations coefficients for the nestedness fraction of beta diversity of the Jaccard dissimilarity index for the taxonomic groups and the environmental distances calculated using EBIC graphical lasso

| | **SP** | **VS** | **UA** | **SFU** | **SB** | **HP** | **SFA** | **TR** |
|---|---|---|---|---|---|---|---|---|
| Soil phosphorus (SP) | 0 | | | | | | | |
| Vegetation structure (VS) | 0 | 0 | | | | | | |
| Understorey arthropods (UA) | -0.10 | 0 | 0 | | | | | |
| Soil fungi (SFU) | -0.10 | 0 | 0 | 0 | | | | |
| Soil bacteria (SB) | 0 | 0 | 0 | -0.10 | 0 | | | |
| Herbaceous plant (HP) | 0 | -0.10 | 0 | 0 | 0 | 0 | | |
| Soil fauna (SFA) | 0 | 0 | -0.10 | 0.11 | 0 | 0 | 0 | |
| Trees (TR) | 0 | 0 | 0 | 0 | 0 | 0 | 0 | 0 |

**Supplementary Table S9.** Partial correlations coefficients for the total beta diversity of the Jaccard dissimilarity index for the functional groups and the environmental distances calculated using EBIC graphical lasso

| | SP | VS | PoUA | PaUA | PrUA | PaFU | SaFU | SyFU | SB | HP | SFA | TR |
|---|---|---|---|---|---|---|---|---|---|---|---|---|
| Soil phosphorus (SP) | 0 | | | | | | | | | | | |
| Vegetation structure (VS) | -0.11 | 0 | | | | | | | | | | |
| Pollinator understorey arthropods (PoUA) | 0 | 0 | 0 | | | | | | | | | |
| Parasitic understorey arthropods (PaUA) | 0 | 0 | 0 | 0 | | | | | | | | |
| Predator understorey arthropods (PrUA) | 0 | 0 | 0 | 0 | 0 | | | | | | | |
| Pathotroph fungi (PaFU) | 0 | 0 | 0 | 0 | 0 | 0 | | | | | | |
| Saprotroph fungi (SaFU) | 0 | 0 | 0 | 0 | 0 | 0.33 | 0 | | | | | |
| Symbiotroph fungi (SyFU) | 0 | 0 | 0 | 0 | 0 | 0.10 | 0.45 | 0 | | | | |
| Soil bacteria (SB) | 0.18 | 0.14 | 0 | 0 | 0 | 0 | 0 | 0.21 | 0 | | | |
| Herbaceous plants (HP) | 0 | 0.14 | 0 | 0 | 0.11 | 0 | 0 | 0 | 0 | 0 | | |
| Soil fauna (SFA) | -0.11 | -0.16 | 0.20 | 0 | 0 | 0 | 0 | 0 | 0 | 0.13 | 0 | |
| Trees (TR) | 0 | 0 | 0.12 | 0 | 0 | 0 | 0 | 0 | 0 | 0 | 0.16 | 0 |

**Supplementary Table S10.** Partial correlations coefficients for the turnover fraction of beta diversity of the Jaccard dissimilarity index for the functional groups and the environmental distances calculated using EBIC graphical lasso

| | **SP** | **VS** | **PoUA** | **PaUA** | **PrUA** | **PaFU** | **SaFU** | **SyFU** | **SB** | **HP** | **SFA** | **TR** |
|---|---|---|---|---|---|---|---|---|---|---|---|---|
| Soil phosphorus (SP) | 0 | | | | | | | | | | | |
| Vegetation structure (VS) | 0 | 0 | | | | | | | | | | |
| Pollinator understorey arthropods (PoUA) | 0 | 0 | 0 | | | | | | | | | |
| Parasitic understorey arthropods (PaUA) | 0 | 0 | 0 | 0 | | | | | | | | |
| Predator understorey arthropods (PrUA) | 0 | 0 | 0 | 0 | 0 | | | | | | | |
| Pathotroph fungi (PaFU) | 0 | 0 | 0 | 0 | 0 | 0 | | | | | | |
| Saprotroph fungi (SaFU) | 0 | 0 | 0 | 0 | 0 | 0.31 | 0 | | | | | |
| Symbiotroph fungi (SyFU) | 0 | 0 | 0 | 0 | 0 | 0.19 | 0.21 | 0 | | | | |
| Soil bacteria (SB) | 0.18 | 0.15 | 0 | 0 | 0 | 0 | 0 | 0 | 0 | | | |
| Herbaceous plants (HP) | 0 | 0.14 | -0.11 | 0 | 0 | 0 | 0 | 0 | 0 | 0 | | |
| Soil fauna (SFA) | 0 | 0 | 0.13 | 0 | 0 | 0 | 0 | 0 | 0 | 0 | 0 | |
| Trees (TR) | 0 | 0 | -0.12 | -0.11 | 0 | 0 | 0 | 0 | 0 | 0.11 | 0 | 0 |

**Supplementary Table S11.** Partial correlations coefficients for the nestedness fraction of beta diversity of the Jaccard dissimilarity index for the functional groups and the environmental distances calculated using EBIC graphical lasso

| | **SP** | **VS** | **PoUA** | **PaUA** | **PrUA** | **PaFU** | **SaFU** | **SyFU** | **SB** | **HP** | **SFA** | **TR** |
|---|---|---|---|---|---|---|---|---|---|---|---|---|
| Soil phosphorus (SP) | 0 | | | | | | | | | | | |
| Vegetation structure (VS) | 0 | 0 | | | | | | | | | | |
| Pollinator understorey arthropods (PoUA) | 0 | 0 | 0 | | | | | | | | | |
| Parasitic understorey arthropods (PaUA) | 0 | 0 | 0 | 0 | | | | | | | | |
| Predator understorey arthropods (PrUA) | 0 | 0 | 0 | 0 | 0 | | | | | | | |
| Pathotroph fungi (PaFU) | 0 | 0 | 0 | 0 | 0 | 0 | | | | | | |
| Saprotroph fungi (SaFU) | 0 | 0 | 0 | 0 | 0 | 0.37 | 0 | | | | | |
| Symbiotroph fungi (SyFU) | 0 | 0 | 0 | 0 | 0 | 0.17 | 0 | 0 | | | | |
| Soil bacteria (SB) | 0 | | 0 | 0 | 0 | 0 | 0 | 0 | 0 | | | |
| Herbaceous plants (HP) | 0 | 0 | 0 | 0 | 0 | 0 | 0 | 0 | 0 | 0 | | |
| Soil fauna (SFA) | 0 | 0 | 0 | 0 | 0 | 0 | 0 | 0 | 0 | 0 | 0 | |
| Trees (TR) | 0 | 0 | -0.12 | 0 | 0 | 0 | 0 | 0 | 0 | 0 | 0 | 0 |

### Supplementary figures

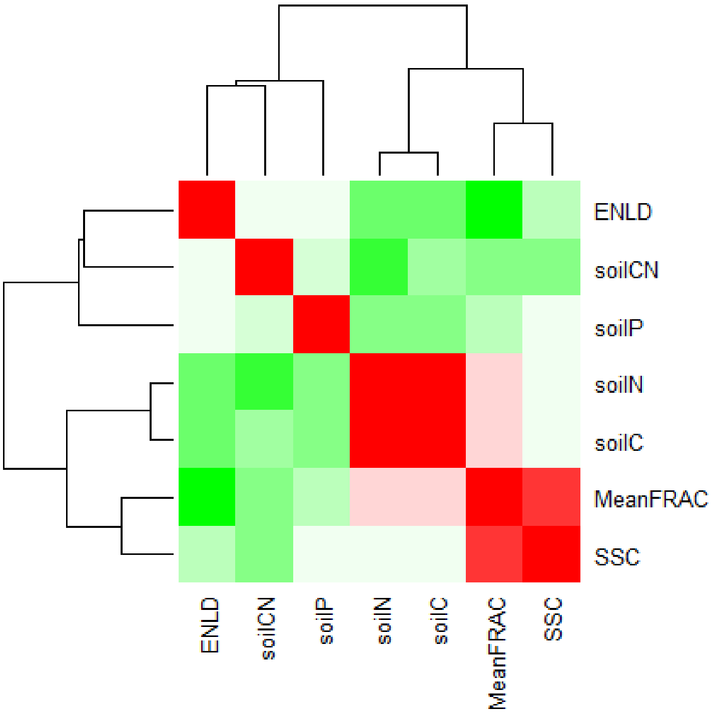


**Supplementary Figure S1.** Correlation matrix among vegetation structural complexity measures and soil nutrient conditions. Red colour means high correlation, white colour means medium correlation, and green colour means low correlation.

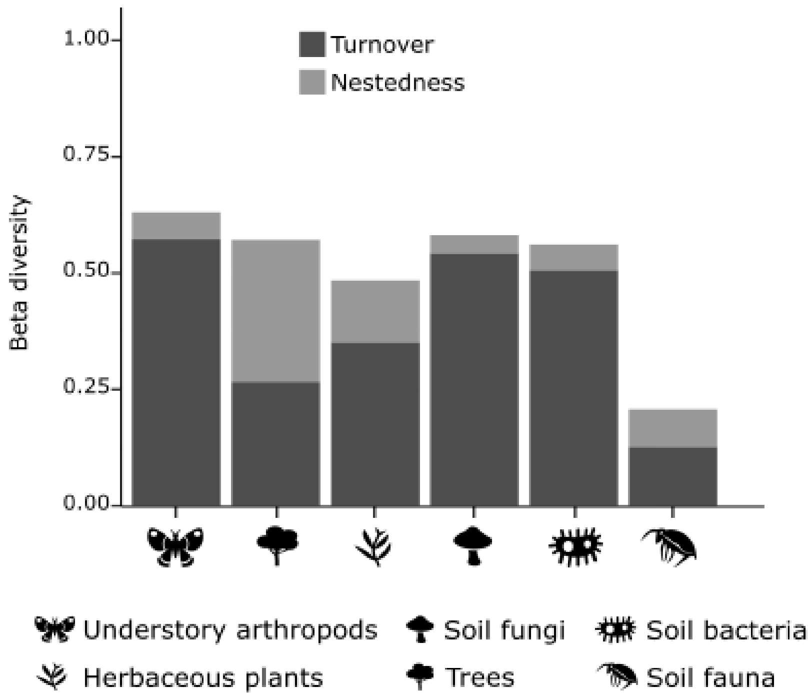


**Supplementary Figure S2. Turnover and nestedness components of beta diversity** for taxonomic groups calculated with Sørensen index.

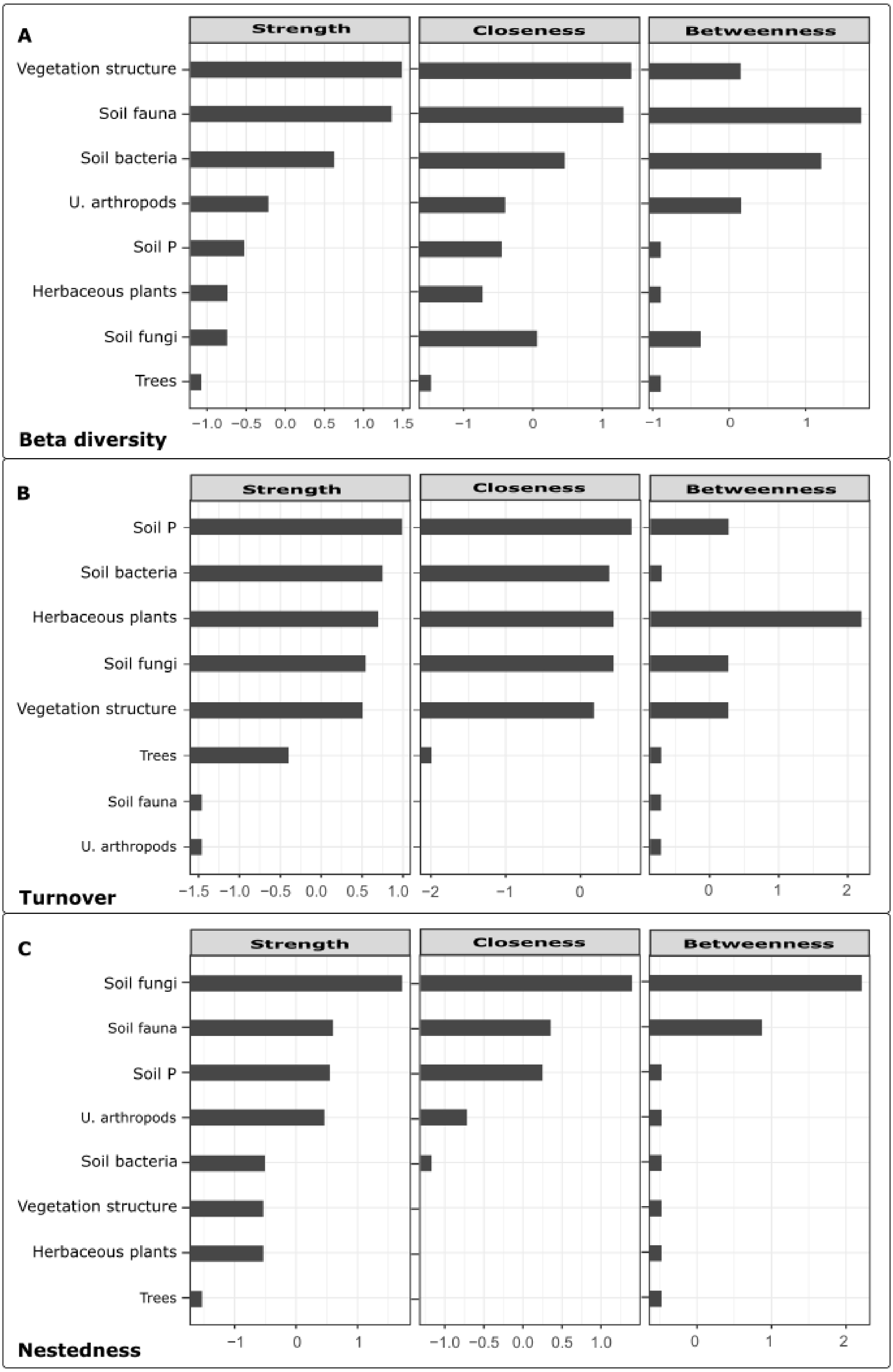
A
Strength
Closeness
Betweenness
Vegetation structure
Soil fauna
Soil bacteria
U. arthropods
Soil P
Herbaceous plants
Soil fungi
Trees
−1.0 −0.5 0.0 0.5 1.0 1.5
−1 0 1
−1 0 1
Beta diversity
B
Strength
Closeness
Betweenness
Soil P
Soil bacteria
Herbaceous plants
Soil fungi
Vegetation structure
Trees
Soil fauna
U. arthropods
−1.5 −1.0 −0.5 0.0 0.5 1.0
−2 −1 0
0 1 2
Turnover
C
Strength
Closeness
Betweenness
Soil fungi
Soil fauna
Soil P
U. arthropods
Soil bacteria
Vegetation structure
Herbaceous plants
Trees
−1 0 1
−1.0 −0.5 0.0 0.5 1.0
0 1 2
Nestedness

**Supplementary Figure S3.** Strength, closeness and betweenness for beta diversity (A), turnover (B), and nestedness (C) for the taxonomic groups and the environmental distances. Centrality indices are shown as standardized z-scores. Low values of correlation stability coefficients (CS (cor = 0.7)) indicate that results should be interpreted carefully (beta diversity betweenness = 0.21, closeness = 0.28, and strength = 0.36. Turnover betweenness = 0, closeness = 0, and strength = 0.44. Nestedness betweenness = 0, closeness = 0, and strength = 0). Strength is correlated with closeness ($R^2$ = 0.88, 0.94, 0.87 for beta diversity, turnover and nestedness, respectively) and with betweenness ($R^2$ = 0.65, 0.31, 0.58 for beta diversity, turnover and nestedness, respectively).

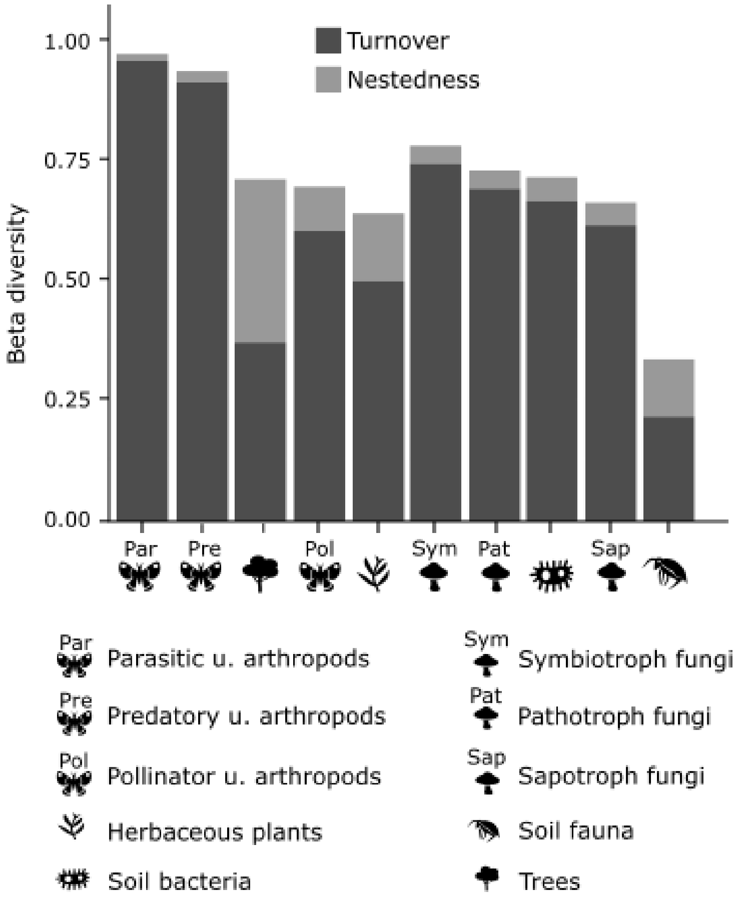


**Supplementary Figure S4. Turnover and nestedness components of beta diversity** for functional groups calculated with Jaccard index. Similar results were found when beta diversity was calculated using Sørensen pairwise dissimilarity (Fig S5).

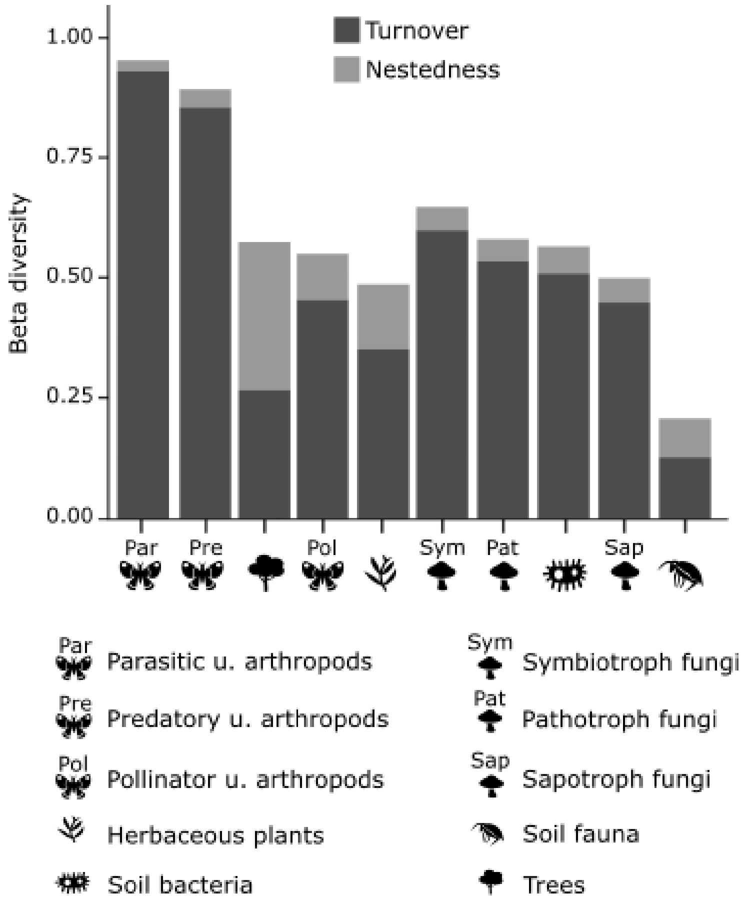


**Supplementary Figure S5. Turnover and nestedness components of beta diversity** for functional groups calculated with Sørensen index.

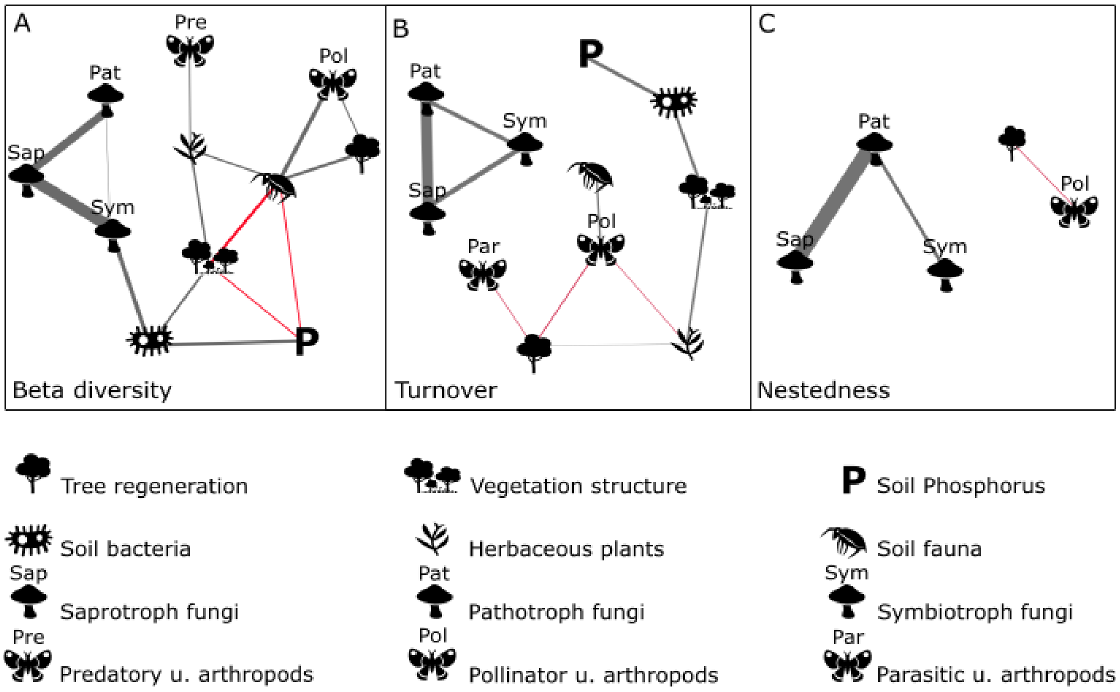


**Supplementary Figure S6**. **The role of landscape heterogeneity, biotic associations, and soil phosphorus in shaping multi-taxa and functional groups beta diversity.** Undirected partial correlation networks between total beta diversity (A), turnover (B), and nestedness (C). Edges thicknesses, i.e., line thickness, are proportional to partial correlation coefficients, with grey and red edges representing positive and negative correlations, respectively. Edge length is not meaningful. The beta diversity network does not include parasitic understorey arthropods and the turnover network does not include predatory understorey arthropods (partial correlation coefficients = 0). The nestedness network only includes soil fungi functional groups, pollinator understorey arthropods and trees (partial correlation coefficients > 0).

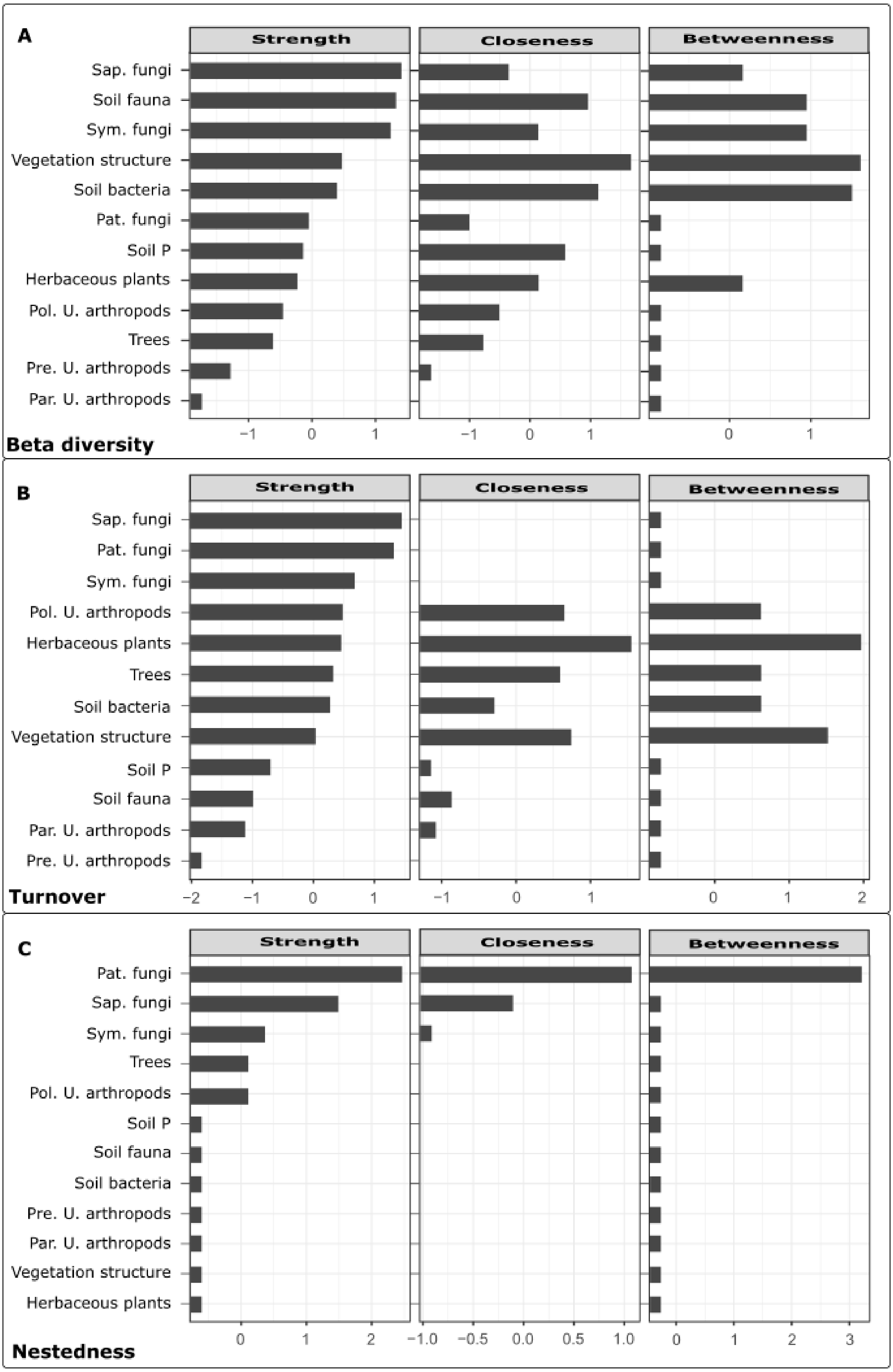

A
Strength
Closeness
Betweenness
Sap. fungi
Soil fauna
Sym. fungi
Vegetation structure
Soil bacteria
Pat. fungi
Soil P
Herbaceous plants
Pol. U. arthropods
Trees
Pre. U. arthropods
Par. U. arthropods
−1
0
1
Beta diversity
B
Strength
Closeness
Betweenness
Sap. fungi
Pat. fungi
Sym. fungi
Pol. U. arthropods
Herbaceous plants
Trees
Soil bacteria
Vegetation structure
Soil P
Soil fauna
Par. U. arthropods
Pre. U. arthropods
−2
−1
0
1
2
Turnover
C
Strength
Closeness
Betweenness
Pat. fungi
Sap. fungi
Sym. fungi
Trees
Pol. U. arthropods
Soil P
Soil fauna
Soil bacteria
Pre. U. arthropods
Par. U. arthropods
Vegetation structure
Herbaceous plants
0
1
2
3
−1.0
−0.5
0.0
0.5
1.0
Nestedness

**Supplementary Figure S7.** Strength, closeness and betweenness for beta diversity (A), turnover (B), and nestedness (C) for the functional groups and the environmental distances. Centrality indices are shown as standardized z-scores. Low values of correlation stability coefficients (CS (cor = 0.7)) indicate that results should be interpreted carefully (beta diversity betweenness = 0.05, closeness = 0, and strength = 0.52. Turnover betweenness = 0, closeness = 0, and strength = 0.28. Nestedness betweenness = 0.44, closeness = 0, and strength = 0.75). Strength is correlated with closeness ($R^2$ = 0.31, 0.65, 0.98 for beta diversity, turnover and nestedness, respectively) and with betweenness ($R^2$ = 0.48, 0.03, 0.59 for beta diversity, turnover and nestedness, respectively).

**Statistical accuracy analysis of the weight of the edges in the network for the taxonomic groups**

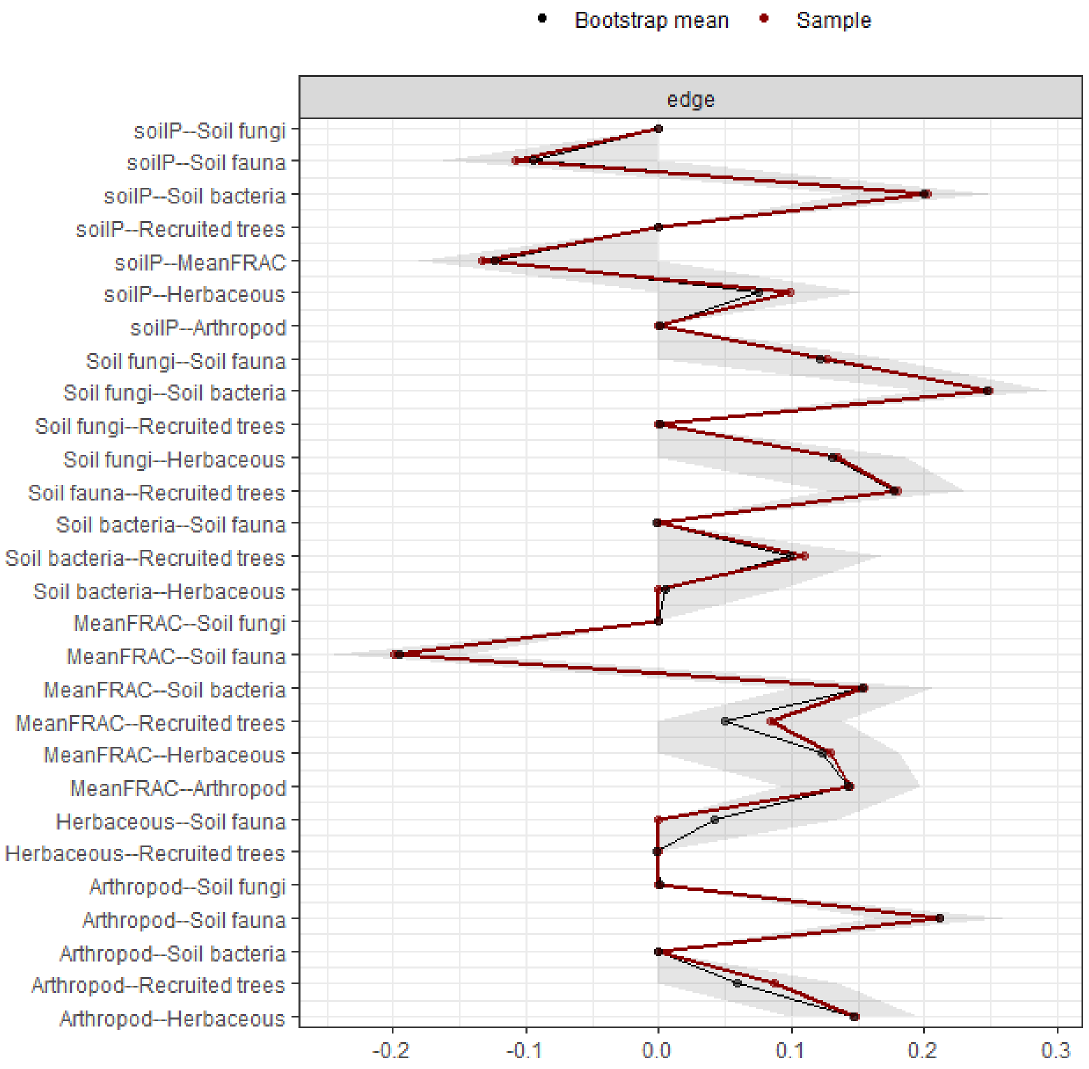


**Supplementary Figure S8.** Observed (red line) and non-parametric bootstrap mean estimated (black line) for the weight of the edges between the vegetation structure complexity, beta diversity across taxonomic groups and soil conditions. Bootstrapped CI: 95% (grey area). The beta diversity was calculated with Jaccard dissimilarity index. MeanFRAC – vegetation structural complexity; Herbaceous – herbaceous plants; Recruited trees – trees; Arthropod – Understorey arthropods.

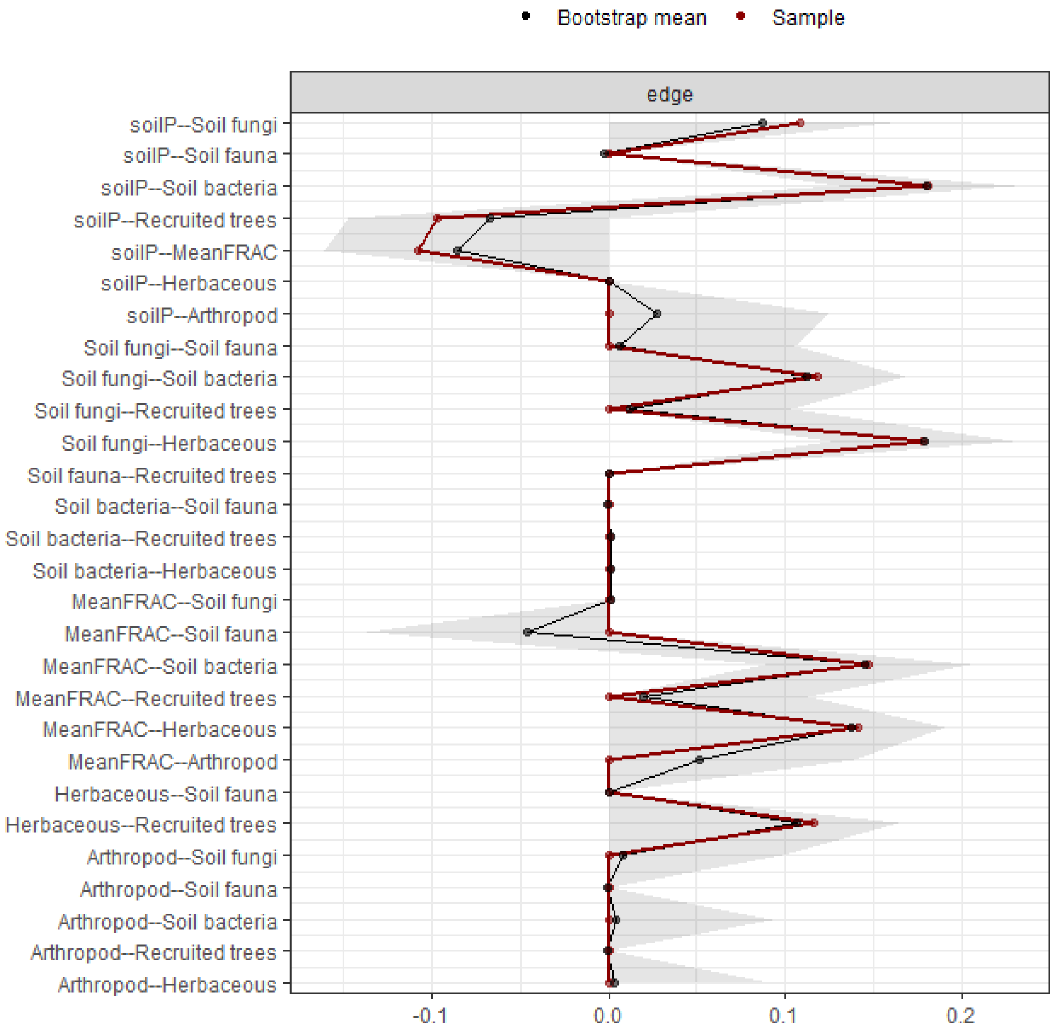


**Supplementary Figure S9.** Observed (red line) and non-parametric bootstrap mean estimated (black line) for the weight of the edges between vegetation structure complexity, turnover across taxonomic groups and soil conditions. Bootstrapped CI: 95% (grey area). The turnover was calculated with Jaccard dissimilarity index. MeanFRAC – vegetation structural complexity; Herbaceous – herbaceous plants; Recruited trees – trees; Arthropod – Understorey arthropods.

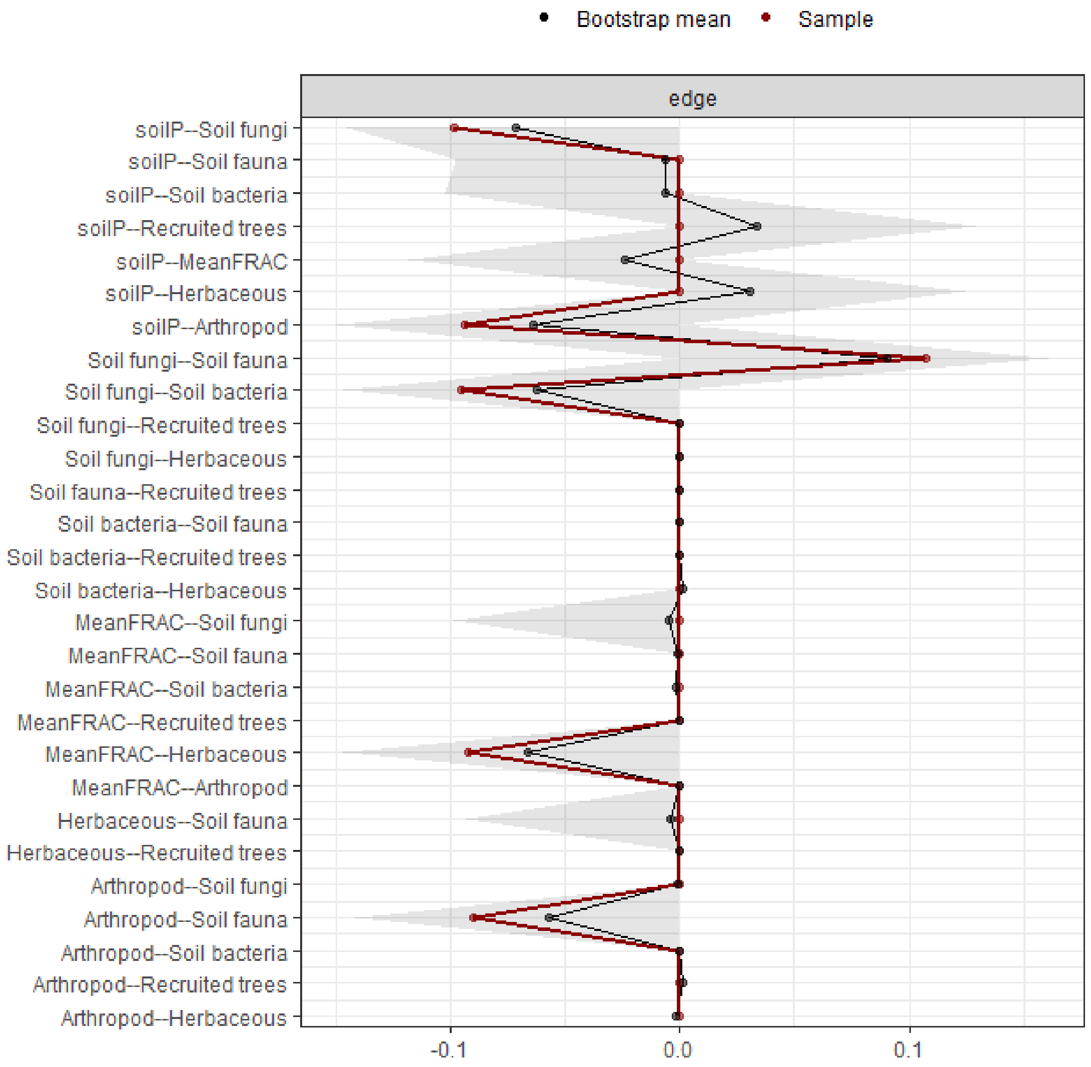


**Supplementary Figure S10.** Observed (red line) and non-parametric bootstrap mean estimated (black line) for the weight of the edges between vegetation structure complexity, nestedness across taxonomic groups and soil conditions. Bootstrapped CI: 95% (grey area). The nestedness was calculated with Jaccard dissimilarity index. MeanFRAC – vegetation structural complexity; Herbaceous – herbaceous plants; Recruited trees – trees; Arthropod – Understorey arthropods

**Statistical accuracy analysis of the weight of the edges in the network for the functional groups**

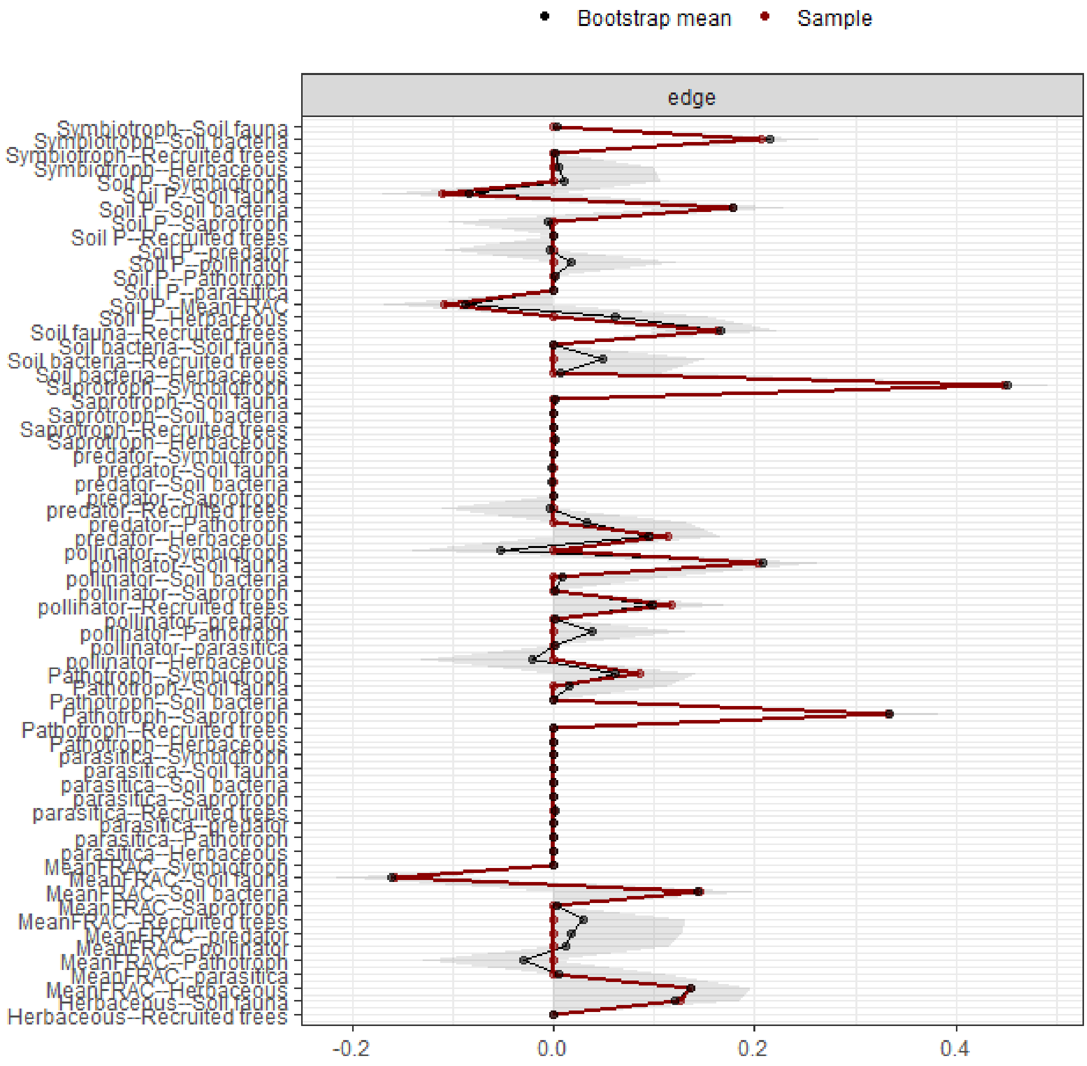


**Supplementary Figure S11.** Observed (red line) and non-parametric bootstrap mean estimated (black line) for the weight of the edges between vegetation structure complexity, beta diversity across taxonomic groups and functional groups, and soil conditions. Bootstrapped CI: 95% (grey area). The beta diversity was calculated with Jaccard dissimilarity index. MeanFRAC – vegetation structural complexity; Herbaceous – herbaceous plants; Recruited trees – trees; Arthropod – Understorey arthropods

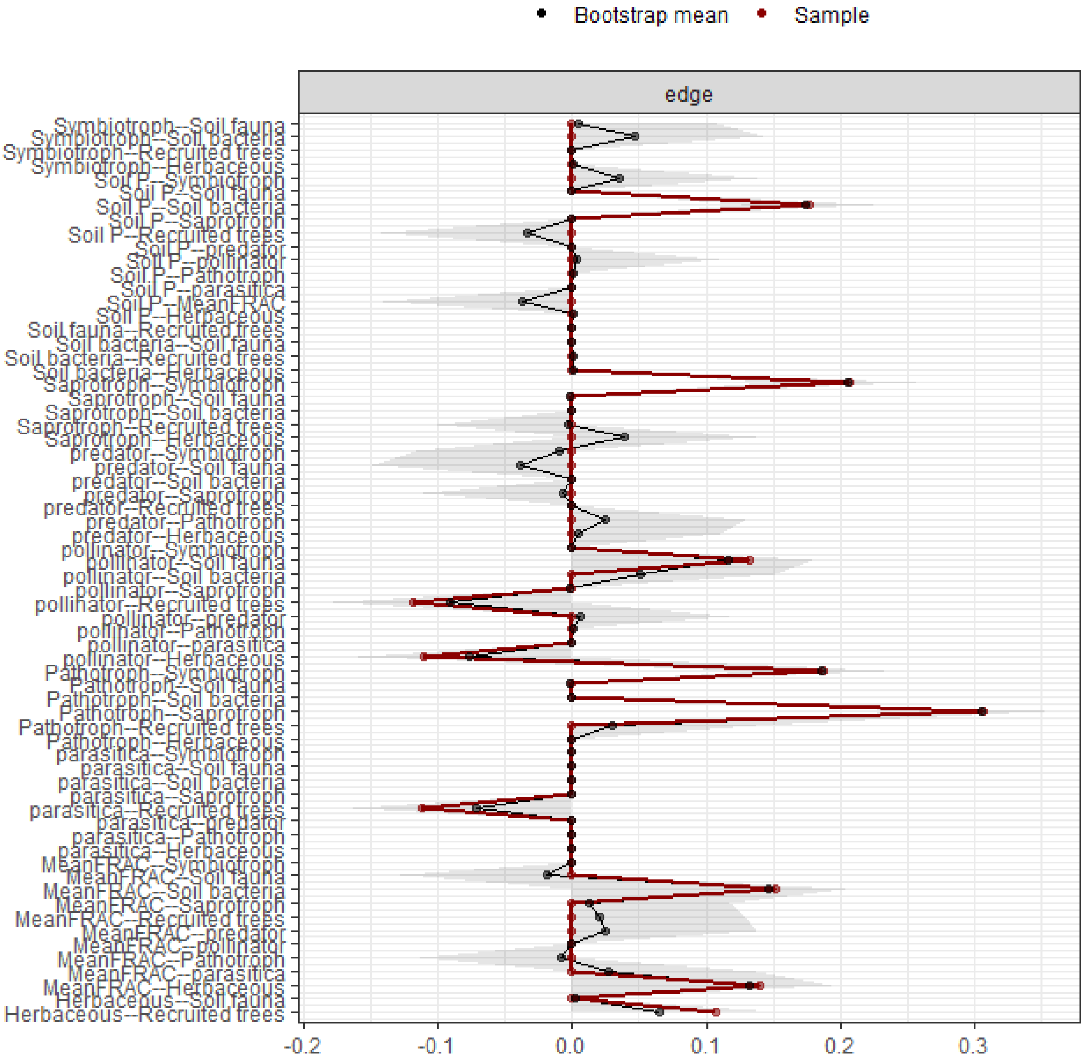


**Supplementary Figure S12.** Observed (red line) and non-parametric bootstrap mean estimated (black line) for the weight of the edges between vegetation structure complexity, turnover across taxonomic groups and functional groups, and soil conditions. Bootstrapped CI: 95% (grey area). The turnover calculated with Jaccard dissimilarity index. MeanFRAC – vegetation structural complexity; Herbaceous – herbaceous plants; Recruited trees – trees; Arthropod – Understorey arthropods

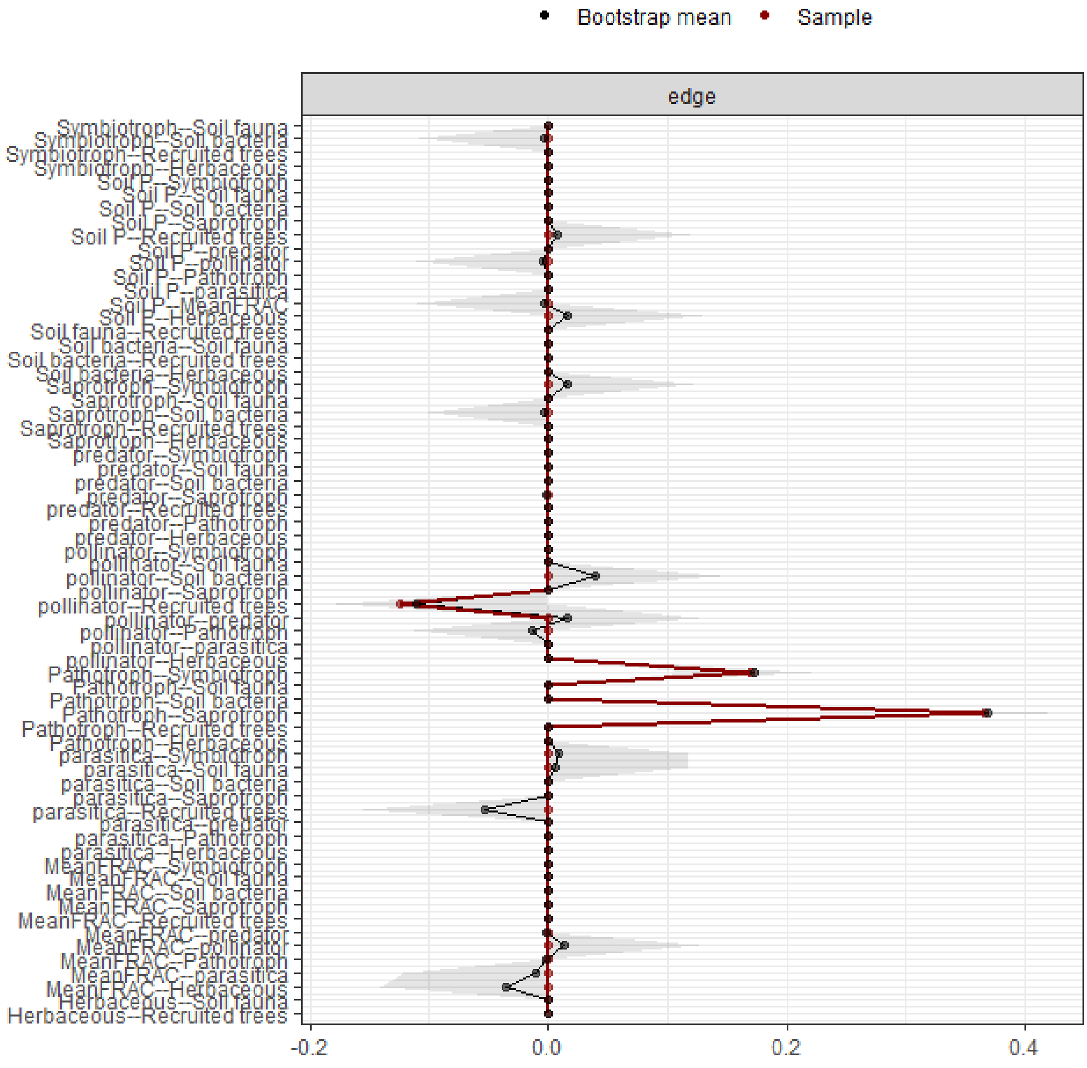


**Supplementary Figure S13.** Observed (red line) and non-parametric bootstrap mean estimated (black line) for the weight of the edges between vegetation structure complexity, nestedness across taxonomic groups and functional groups, and soil conditions. Bootstrapped CI: 95% (grey area). The nestedness was calculated with Jaccard dissimilarity index. MeanFRAC – vegetation structural complexity; Herbaceous – herbaceous plants; Recruited trees – trees; Arthropod – Understorey arthropods.
